# The effect of DNA methylation on DNA structure and DNA-small molecule interactions

**DOI:** 10.64898/2026.07.30.741680

**Authors:** Gözdem Çavdar, Nurdan Emin, Aybüke Gülkaya, Deniz İrem Alpinanç, Antoine Marion, Özgül Persil Çetinkol

## Abstract

The methylation of cytosines at the 5^th^ position (d5mC) is one of the most common epigenetic modifications, and alterations in methylation profile of cells are known to be involved in progression of many diseases including cancer. Increased stability of DNA accompanied by decreased flexibility upon methylation is thought to be a reason behind methylation profiles. The effects of d5mC on DNA stability and structure were investigated via systematic changes in the number and position of d5mCs in DDD. Our results revealed that d5mC substitutions changed DNA conformation and increased the stability only slightly. Next, the effect of DNA methylation on DNA-small molecule interactions was investigated using DDD and fully methylated analogue, DDD8. All the molecules examined (EtBr, Dox, Net and Hoe) had slightly higher affinity to DDD8 compared to DDD. Conversely, their effect, especially Dox’s, on DDD structure was more pronounced. Further investigations via MD simulations revealed high selectivity of Dox towards a single intercalation site where the methoxy group of Dox interacts with the methyl group of d5mC and that of two precedent dT to create a highly stable hydrophobic cluster. Hydrophobic cluster formation was not observed upon Dox binding to DDD. Our results rationalize the increased stability of DDD8 over DDD, and open new routes for the design of drugs targeting epigenetic modifications. We suggest, the design of drugs that can engage in hydrophobic interactions with methyl groups in the major groove of a 5’-dTdTd5mCdG-3’ sequence might lead the way in specific targeting of hypermethylated regions in cancer cells.

## INTRODUCTION

In eukaryotes, to fit within the cell, DNA is packaged around histone proteins forming nucleosomes that are the basic repeating units of chromatin.^1^ This higher level of organization is known to regulate many cellular processes such as gene expression, chromosome segregation, and DNA replication just by adjusting the DNA accessibility.^2^ The DNA accessibility, thus the chromatin structure, is known to be mainly regulated by epigenetic modifications in DNA and histones.^3^

In mammals, the main type of epigenetic modification, DNA methylation, occurs by the transfer of a methyl group from S-adenosyl methionine to the fifth carbon of a cytosine base to produce 5-methylcytosine (5mC) by DNA methyltransferases (DNMTs). This reaction is known to be taking place mainly in CpG islands^4^ and the presence of methyl groups on CpG islands of the promoter regions are believed to be modulating gene expression by decreasing the flexibility and curvature of DNA as a result of steric effect and hydrophobicity.^5,6^

To reveal the exact role of DNA methylation on DNA structure and stability, studies involving the substitution of one cytidine (dC) with 5-methylcytidine (d5mC) in Dickerson dodecamer (DDD, 5’-CGCGAATTCGCG-3’) **(Table 1)** were performed. A slight increase in the thermal denaturation temperatures of DDDs was observed upon the substitution of only one dC with d5mC.^7,8,9^ Substitution of more than one dC with d5mC was also studied by Theruvathu et al. using DDD.^10^ They investigated the effect of DNA methylation on DNA structure and stability by substituting the 3^rd^, the 9^th^ or both of the 3^rd^ and the 9^th^ dCs. Contrary to previous studies, they found no differences in either thermal or thermodynamic stability between the substituted and unsubstituted dsDNAs. They also established via crystallography and ^1^H NMR that the dsDNA containing d5mC only at the 3^rd^ position was a standard B-form DNA and was not significantly different from the unsubstituted duplex. In addition, Tsuruta et. al. investigated the effect of DNA methylation on 10-mer duplexes and reported a relatively significant stabilization, up to 8°C, upon methylation.^11^ The role of methylation on DNA structure was investigated to a great extend via computational studies.^12,13,14,15^ Hognon et.al. determined the effect of different levels of methylation in CpG islands on the structural parameters of dsDNA made of 20 base pairs.^16^ The authors concluded that local methylation has little to no effect on the DNA structure, while a high density of d5mC in CpG islands induces pronounced structural changes, mainly affecting the zeta-angle and major groove accessibility.

**Table 1.** DNA sequences (Dickerson Dodecamer) used in the experiments.

| Short Name for the Sequence | DNA Sequence | DNA Duplexes |
| --- | --- | --- |
| DDD | 5'-CGCGAATTCGCG-3' | 5'-CGCGAATTCGCG-3'<br>3'-GCGCTTAAGCGC-5' |
| DDD2a | 5'-<sup>me</sup>CGCGAATTCGCG-3' | 5'-<sup>me</sup>CGCGAATTCGCG-3'<br>3'-GCGCTTAAGCG<sup>me</sup>C-5' |
| DDD2b | 5'-CG<sup>me</sup>CGAATTCGCG-3' | 5'-CG<sup>me</sup>CGAATTCGCG-3'<br>3'-GCGCTTAAG<sup>me</sup>CGC-5' |
| DDD4a | 5'-<sup>me</sup>CG<sup>me</sup>CGAATTCGCG-3' | 5'-<sup>me</sup>CG<sup>me</sup>CGAATTCGCG-3'<br>3'-GCGCTTAAG<sup>me</sup>CG<sup>me</sup>C-5' |
| DDD4b | 5'-<sup>me</sup>CGCGAATTCG<sup>me</sup>CG-3' | 5'-<sup>me</sup>CGCGAATTCG<sup>me</sup>CG-3'<br>3'-G<sup>me</sup>CGCTTAAGCG<sup>me</sup>C-5' |
| DDD4c | 5'-CG<sup>me</sup>CGAATT<sup>me</sup>CGCG-3' | 5'-CG<sup>me</sup>CGAATT<sup>me</sup>CGCG-3'<br>3'-GCG<sup>me</sup>CTTAAG<sup>me</sup>CGC-5' |
| DDD6a | 5'-<sup>me</sup>CG<sup>me</sup>CGAATT<sup>me</sup>CGCG-3' | 5'-<sup>me</sup>CG<sup>me</sup>CGAATT<sup>me</sup>CGCG-3'<br>3'-GCG<sup>me</sup>CTTAAG<sup>me</sup>CG<sup>me</sup>C-5' |
| DDD6b | 5'-<sup>me</sup>CG<sup>me</sup>CGAATTCG<sup>me</sup>CG-3' | 5'-<sup>me</sup>CG<sup>me</sup>CGAATTCG<sup>me</sup>CG-3'<br>3'-G<sup>me</sup>CGCTTAAG<sup>me</sup>CG<sup>me</sup>C-5' |
| DDD8 | 5'-<sup>me</sup>CG<sup>me</sup>CGAATT<sup>me</sup>CG<sup>me</sup>CG-3' | 5'-<sup>me</sup>CG<sup>me</sup>CGAATT<sup>me</sup>CG<sup>me</sup>CG-3'<br>3'-G<sup>me</sup>CG<sup>me</sup>CTTAAG<sup>me</sup>CG<sup>me</sup>C-5' |

DNA hypomethylation and site-specific promoter hypermethylation are known to play roles in certain cancers due to the induced chromosome instability and transcriptional repression of tumor suppressor genes, respectively.^17^ Targeting these hypomethylation and hypermethylation sites specifically using chemotherapeutic agents is one of the emphasized areas of research in the last decade.^18,19^ Altering the methylation levels via chemotherapeutic agents is considered to be a route to alter many cellular processes. For instance, inhibitors of both histone methyltransferases (HMTs) and DNA methyltransferases (DNMT) are shown to reduce metastasis.^18^ Many small molecule chemotherapeutic agents such as doxorubicin, mitoxantrone, actinomycin are currently used in chemotherapy to target the DNA structure and its biological functions.^20^ However, these agents are specific to neither hypermethylated nor hypomethylated sequences. To design small molecules that are specific to hypermethylated or hypomethylated sites, an understanding of the effect of methylation on the binding dynamics of these molecules to DNA is necessary.

Accordingly, we investigated the role of DNA methylation on DNA structure and stability by systematically changing the number and the placement of 5-methylcytidines in a model Dickerson dodecamer dsDNA. In addition, we investigated the effect of DNA methylation on DNA-small molecule interactions. Intercalators act by entering between two adjacent base pairs, resulting in unwinding and extension of DNA. Such a structural change is known to prevent the recognition of DNA by DNA-associated proteins. On the other hand, minor groove binders that are known to bind preferably to AT-rich regions, and interfere with the binding of other macromolecules such as proteins and enzymes through the grooves to DNA, thus with the cell cycle.^20^ We chose ethidium bromide (EtBr) and doxorubicin (Dox) as model intercalators, hoechst 33258 (Hoe) and netropsin (Net) as model minor groove binders. Our results revealed that even though DNA methylation increases the stability and changes the conformation of DNA only slightly without altering the standard B-form conformation, it has profound effects on small molecule binding. Further molecular dynamics investigations on Dox binding to DDD8 revealed the formation of a hydrophobic cluster that plausibly drives the binding of Dox to a specific intercalation site. Neither the hydrophobic cluster formation nor the specificity was observed upon Dox binding to DDD. We believe that designing small molecules with the ability to form such hydrophobic pockets, might help to regulate gene expression in specific hypermethylated sites that are associated with certain cancer types.

## MATERIALS AND METHODS

Oligonucleotides **(Table 1)** were purchased from Integrated DNA Technologies (Europe) in a lyophilized form and used without further purification. Stock solutions of the oligonucleotides were prepared by dissolving the solid in millipore water. The concentrations of stock solutions were determined from the UV-Vis absorbances using the extinction coefficients supplied by the manufacturer at 260 nm: DDD 110700 M^-1^cm^-1^; DDD2a and DDD2b 109000 M^-1^cm^-1^; DDD4a, DDD4b and DDD4c 107300 M^- 1^cm^-1^; DDD6a and DDD6b 105600 M^-1^cm^-1^; DDD8 103900 M^-1^cm^-1^. EtBr, Net, and Hoe were purchased from Sigma and used without further purification. Dox was obtained from Zhejiang Hisun Pharmaceutical CO., LTD (Zhejiang, China). Stock solutions of small molecules were also prepared by dissolving weighed amounts in millipore water and stored in the dark at 4°C. Concentrations of stock solutions were determined by UV-Vis spectroscopy using the following extinction coefficients: EtBr ɛ_478_ 5680 M^-1^cm^-1^,^37^ Dox ε_480_ 11500 M^-1^cm^-1^,^38^ Net ε_296_ 21500 M^-1^cm^-1^,^39^ and Hoe ε_338_ 42000 M^-1^cm^-1^.^40^ All the experiments were carried out using BPES buffer (pH = 7.0) including 6 mM Na_2_HPO_4_, 2 mM NaH_2_PO_4_, 1 mM Na_2_EDTA, and 185 mM NaCl.

In both UV-Vis and CD experiments, the ratio of DNA to intercalators, EtBr or Dox, was kept as 4 DNA bases:1 small molecule in accordance with the ‘nearest neighbor exclusion’ principle.^41^ The ratio of DNA to groove binders, Net and Hoe was kept at 1 DNA duplex:1 small molecule in accordance with the crystal structures.^42,43^ The samples contained 60 µM of DNA in base and/or a preferred small molecule in an amount according to the rules explained. For the formation of the proper duplex, before the experiments, each oligonucleotide was heated up to 95°C in a water bath, kept at this temperature for 5 minutes, and let cool down to room temperature for overnight. The preparation of samples including both DNA and a small molecule was applied by adding the small molecule to the solution including annealed DNA.

### Circular Dichroism Spectroscopy

The spectra of samples were collected by JASCO J-815 Circular Dichroism (CD) spectrophotometer coupled with a Peltier temperature control unit (PTC-423S/15), between 215 and 500 nm. For thermal denaturation experiments, the samples were heated from 5 to 95°C with 2°C intervals, 200 nm/min scanning speed and 1 nm band width. The samples were kept at the same temperature for 2 minutes before the collection of the spectrum. Thermal denaturation curves were obtained by monitoring the change in ellipticity at 250 nm with respect to temperature.

### UV-Visible Spectroscopy

JASCO V-730 UV-visible spectrophotometer coupled with ETCS-761 temperature control panel was used in UV-Vis experiments. A programmable peltier cell holder was used in thermal denaturation experiments. Both heating and cooling thermal denaturation profiles were obtained between 15°C and 85°C at each temperature with 1°C per minute heating rate. The experiment was repeated twice and the UV-Vis absorbance at 260 nm was monitored for these two experiments. Then, thermal denaturation curves were obtained by the average of absorbance values with respect to temperature and the melting temperatures were obtained from the middle points of normalized curves.

### Fluorescence Spectroscopy

The samples including 1.0 µM of the corresponding small molecule were titrated with the sample containing 1.0 µM of the same molecule and 400 µM (in base pair) DDD or DDD8. After each titration, emission spectrum was collected by Agilent Cary Eclipse Fluorescence spectrophotometer, by using excitation wavelengths of 510 nm for EtBr, 480 nm for Dox, and 365 nm for Hoe, and emission wavelength of 525 to 800 nm for EtBr, 500 to 750 nm for Dox, and 375 to 675 nm for Hoe. Excitation and emission slits were set to 10.0 and 10.0 for EtBr, 5.0 and 10.0 for Dox, and 2.5 and 5.0 for Hoe, respectively. For Dox and Net, the association constants were obtained by fitting the integrated fluorescence intensity as a function of DNA concentration using the least squares equations as reported previously.^44,45^ For Hoe, the data was fit to sigmoidal curve better, and the association constants were obtained from the mid points of the curves.

### Molecular Modelling

All molecular dynamics simulations were performed with the CUDA implementation of pmemd in Amber18.^46^ Parameters for canonical DNA bases were taken from the BSC1 force field.^47^ Specific point charges were derived for Dox and 5-methyl-cytosine (5mC) using the RESP^48^ methodology from electrostatic potentials calculated at the HF/6-31G* with Orca 4.2.1.^49,50^ by following the standard Amber strategy.^51^ Dox was modelled with its amine group protonated and gaff^52^ parameters were assigned automatically by antechamber as part of the AmberTools19 distribution.^46^ All parameter files are available upon reasonable request to the authors. Initial geometries of wild type Dickerson dodecamer (DDD) and fully methylated (DDD8) were generated by the nucleic acid builder (nab) as part of the AmberTools19 distribution.^46^ Structures of DDD and DDD8 with one Dox molecule intercalated were generated manually with the Pymol software^53^ and further subject to a short MD simulation in the gas phase using a softcore potential^54^ locally implemented in sander, to smoothly let the intercalation site adapt to the presence of the ligand. Considering the symmetry of the system, only six intercalation sites were considered, with two orientations of the daunosamine group of Dox; i.e., either pointing towards the center of the sequence or towards the end (respectively labelled as “inward” and “outward”).

A total of 26 distinct systems were subject to molecular dynamics simulations, i.e., apo DDD and DDD8, as well as 12 intercalated versions for each DNA sequence (6 sites with two orientations of the daunosamine group of Dox). Each system was solvated in a truncated octahedral box of TIP4Pew water molecules^55^ with a buffer region of 12 Å between solute atoms and box edges. Na and Cl ions were added to ionize the system to an approximate concentration in NaCl of 0.5 M^56^ with extra Na ions to neutralize the simulation box. After minimization, each box was gradually heated up to 100 K in the NVT ensemble, and subsequently to 300 K in the NPT ensemble. Inter-bases harmonic distance-restraints were used for intercalated systems until reaching 200 K to preserve Watson-Crick base pairing during heat up. All boxes reached an equilibrated density of about 1.05 g cm^-3^ by the end of the heat up procedure (data not shown). For each system, two production simulation runs were conducted with randomly assigned initial velocities for 750 ns in the NPT ensemble. All MD simulations used periodic boundary conditions with particle mesh Ewald treatment^57,58,59,60^ of long-range electrostatic interactions, a cut off of 12 Å for non-bonded terms in the real space, the SHAKE algorithm to constrain bonds involving hydrogen atoms, and an integration time step of 2 fs. Langevin dynamics^61^ with a collision frequency of 2 ps^-1^ and the Berendsen barostat^62^ were used to control temperature and pressure, respectively.

Interaction energy between Dox and DNA was evaluated over the last 500 ns of each simulation run with the generalized Born/surface area (GBSA) model^63^ using the mbondi3 set of atomic radii and the modified GBn model (i.e., igb = 8).^64^ The contribution of DNA deformations to the total binding energy was evaluated by comparing the energy of apo and intercalated DNA as averaged over the last 500 ns of the corresponding simulations runs (two runs per system) by performing single point calculations on the isolated DNA sequences with the GBSA model from frames extracted every 100 ps.

Analysis was performed with the CURVES+ and Canal codes^65^ as well as with the cpptraj suite^66^ as implemented in AmberTools19. Time series were calculated over the entire simulation time, while the distribution of the various descriptors considered in this work was obtained from the last 500 ns of each simulation run for a given system (i.e., 1 ms total) with frames extracted every 200 ps.

## RESULTS AND DISCUSSION

The effect of the presence of methyl groups on the secondary structure of DNA was first investigated by circular dichroism (CD) spectroscopy. The CD spectrum of the non-methylated Dickerson dodecamer (DDD) recorded at 5°C is in accordance with the previously published CD spectra for DDD and is characterized as the spectrum of B-DNA with a positive band at around 280 nm and a negative band at around 250 nm **(Figure 1)**.^8, 21^

**Figure 1.**
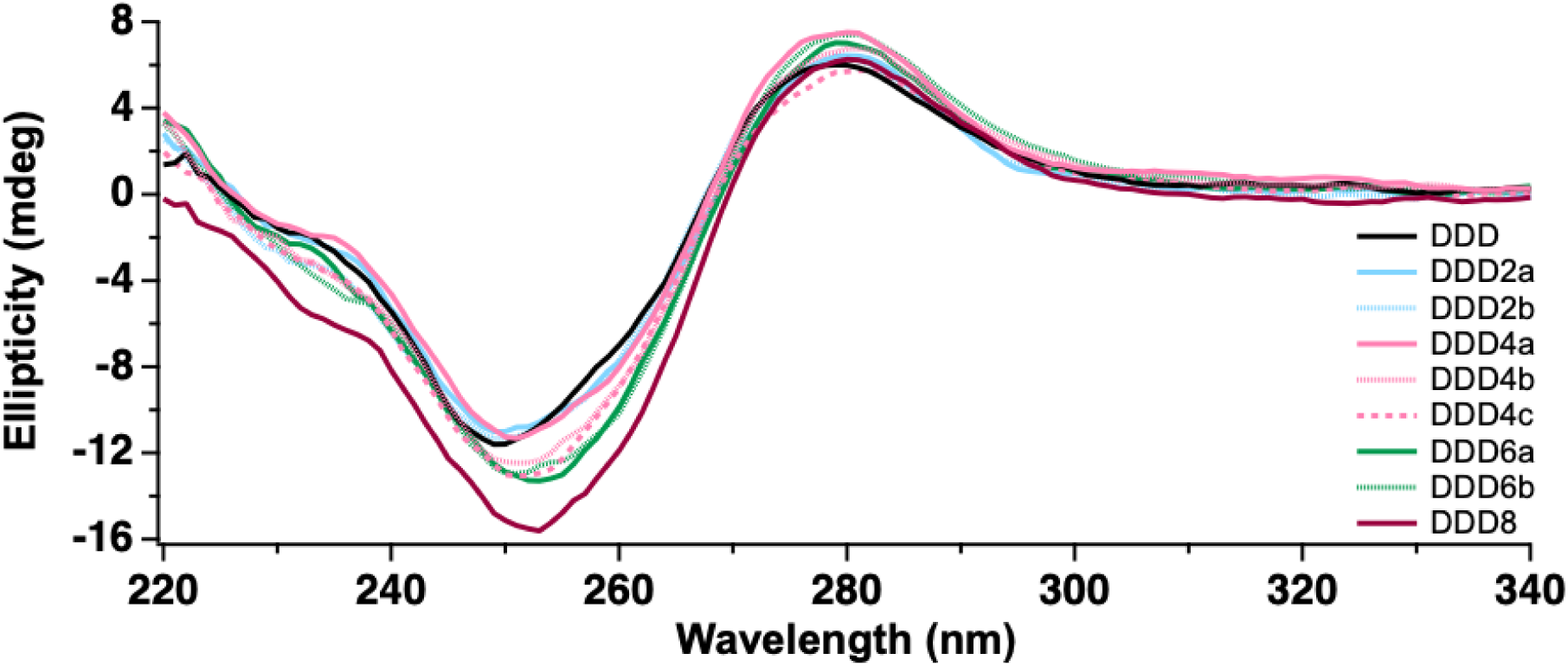
CD spectra of DDD, and d5mC substituted DDDs at 5°C. The sequences are given explicitly in Experimental Section, **Table 1**.

Next, the CD spectra of dodecamer sequences with two (DDD2a and DDD2b; d5mCs vary in position), four (DDD4a, DDD4b and DDD4c), six (DDD6a and DDD6b) or eight (DDD8) d5mCs were acquired and compared to the CD spectrum of DDD. As displayed in **Figure 1**, the presence of several d5mCs did not alter the CD spectrum significantly, indicating that methylation has little to no effect on the structure of DDD, as previously established by Renciuk et al.^8^, Theruvathu et al.^10^, and Tsuruta et al.^11^ While the main features of a B-DNA CD spectrum are preserved upon methylation, we noticed substantial changes in the wavelength and the intensity of the minimum absorption with increasing number of methyl groups. As displayed in **Figure 1**, the CD spectra of DDD, DDD2a, DDD2b and DDD4a are similar to each other. However, the band centered at 250 nm is red shifted by 2 nm and its intensity is decreased from -11.6 mdeg to -13.0 mdeg with increasing methylation levels in DDD4b, DDD4c, DDD6a, and DDD6b. The red-shift and change in intensity are more pronounced in DDD8, where all the dCs are substituted with d5mCs. The spectrum of DDD8 is red shifted by about 4 nm, and the change in intensity is about 4 mdegs, from -11.6 mdeg to -15.6 mdeg, compared to DDD. These results indicate that, even though the d5mC substitution does not alter the helical B form structure, it does affect π-π* transitions in the plane of the bases.^21^ The effect was more pronounced when all the dCs were substituted with d5mCs. However, the change in ellipticity was not linearly correlated with the number of d5mC substitutions. Even though, DDD, DDD2a, DDD2b and DDD4a have different number of d5mCs, the ellipticity values obtained at 250 nm are close to each other. Again, DDD4b, DDD4c, DDD6a and DDD6b have almost the same ellipticity at 250 nm. Moreover, when the CD spectra of DDD’s with the same number of substitutions such as DDD4a, DDD4b and DDD4c are compared, it appears that the location of the substitution does not significantly affect the electronic transitions as does the number of substitutions.

To determine the effect of methyl groups on the thermal denaturation temperatures (T_m_) of DNA duplexes, thermal denaturation experiments were performed. The difference in thermal stability was first investigated with CD spectroscopy, but the thermal denaturation profiles obtained were very similar **(Supplementary Figure S1)**. However, the thermal denaturation profiles obtained via UV-Vis revealed an increase, yet not a linear correlation, in thermal denaturation temperature with the increasing degree of methylation **(Supplementary Figure S2)**. Thermal denaturation temperatures obtained from the normalized profiles are listed in **Table 2**. The greatest increase was observed for the fully methylated sequence (DDD8), where the denaturation temperature is 3°C higher than that of the reference DDD. This difference corroborates earlier observations indicating that the presence of methyl groups increases the stability of the duplex structure **(Table 2, Supplementary Figure S2)**.^7, 8, 9^

**Table 2.**
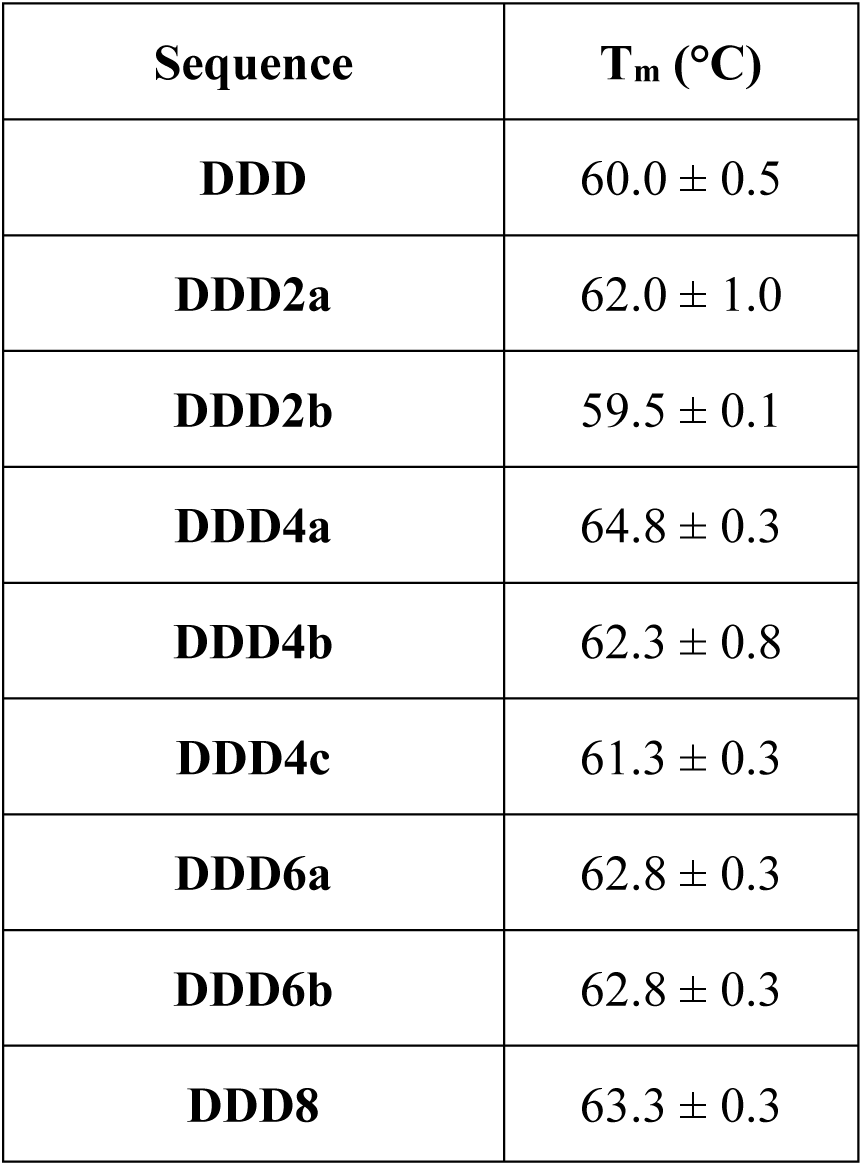
Thermal denaturation temperatures obtained by monitoring the change in UV- Vis absorbance at 260 nm.

To assess the effect of DNA methylation on the molecular recognition of DNA by small molecules, the interactions of DDD and DDD8 with intercalators or minor groove binders was investigated via UV-Vis and CD experiments. **Figure 2** displays the UV-Vis spectra of DDD and DDD8 in the presence and in the absence of small molecules. The change in absorption in the longer-wavelength region of the UV-Vis spectra of all the small molecules upon binding to DDD or DDD8 confirms their interaction with the dsDNA. The absorption maxima of all molecules are shifted to the red upon binding to DNA. We observed a decrease in the intensity of the absorption maximum (hypochromic effect) of the intercalators, EtBr **(Figure 2A)** and Dox (Figure 2B), and an increase for the groove binder Hoe **(Figure 2D)**. Upon the binding of Net to DDD or DDD8, neither hypochromism nor hyperchromism was observed **(Figure 2C).** The hypochromic effect observed for Dox appears to be slightly more pronounced upon binding to DDD8 compared to DDD **(Figure 2B)**. No such difference related to the degree of methylation was observed for EtBr **(Figure 2A)** or Hoe **(Figure 2D)**.

**Figure 2.**
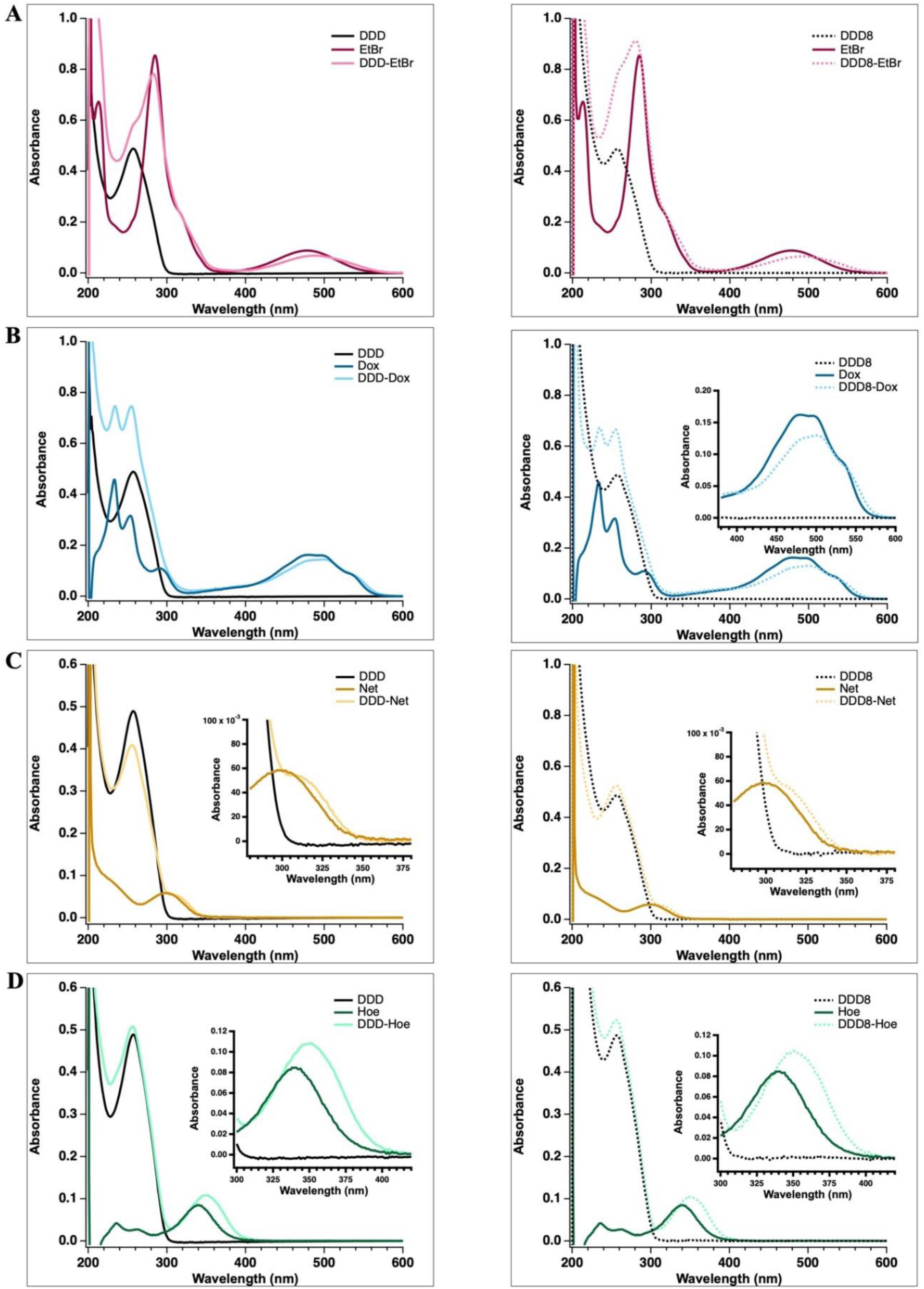
UV-Vis spectra of DDD and DDD8 in the absence and presence of EtBr **(A)**, Dox **(B)**, Net **(C)**, and Hoe **(D)** at 15°C. The insets show the longer wavelength region of the UV-Vis spectra.

**Figure 3** displays the CD spectra of DDD and DDD8 in the presence and in the absence of small molecules. As mentioned above, the CD spectra of DDD and DDD8 in the absence of a small molecule are similar and both correspond to a standard B-form DNA, with a negative peak at around 250 nm and a positive peak at around 280 nm **(Figure 1)**.^8,22^ The main features of the CD spectra mostly change in a similar manner for DDD and DDD8 upon the binding of EtBr, Net and Hoe **(Figure 3A, C, D)**. **Figure 3A** displays the CD spectra of DDD and DDD8 in the presence and absence of EtBr. The intensity of the peak at ∼250 nm was decreased, and the peak at ∼280 nm was shifted to ∼274 nm upon EtBr binding. In addition, the formation of a new local minimum at ∼228 nm and a weak induced band between 300 nm and 370 nm were observed.^23, 24^ The induced CD band which provides evidence for the interaction of achiral EtBr with the chiral environment of DNAs was slightly red-shifted in the spectra of DDD8-EtBr compared to DDD-EtBr.

**Figure 3.**
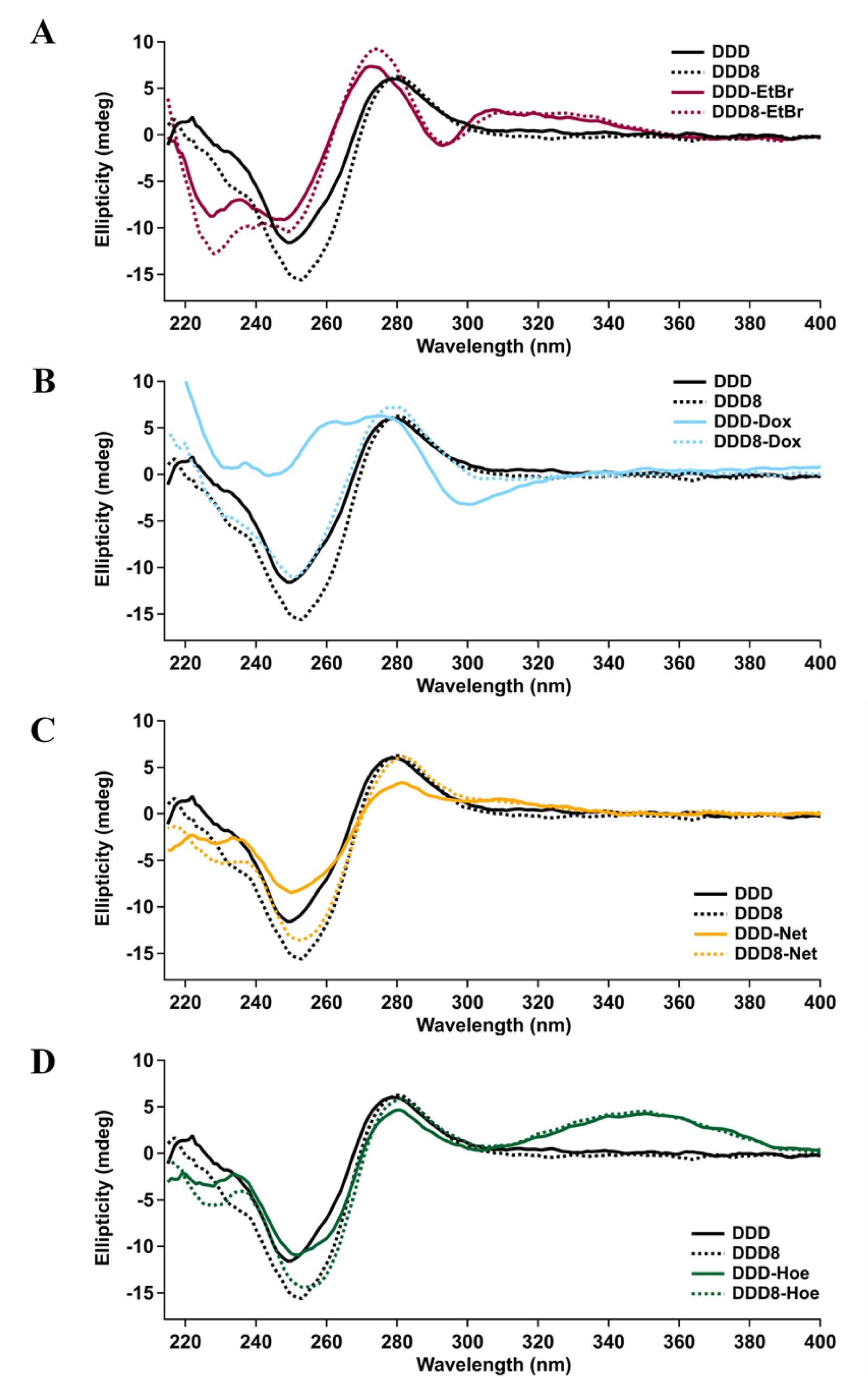
CD spectra of DDD and DDD8 in the absence and presence of EtBr **(A)**, Dox **(B)**, Net **(C)**, and Hoe **(D)** at 15°C.

The effect of Net binding on the secondary structure of DNA is slightly more pronounced in DDD compared to DDD8 **(Figure 3C)**. Nevertheless, the CD spectra of DDD-Net and DDD8-Net are similar to each other shape-wise. The change in the intensity of both bands at ∼250 nm and ∼280 nm is stronger in the spectrum of DDD upon Net binding. Only the intensity of the band at 250 nm is slightly decreased upon binding of Net to DDD8. In other words, no change was observed in the intensity of the band at ∼280 nm. The formation of a very weak induced CD band between 300-340 nm was observed both for DDD-Net and DDD8-Net complexes.

In contrast to the other molecules considered in the present work, the differences in the CD spectra of DDD and DDD8 induced upon Dox binding are substantial **(Figure 3B)**. In DDD, the peak with the local minimum at ∼250 nm is split into two upon Dox binding with local minima at ∼233 nm and ∼243 nm, and a loss of intensity. The peak at ∼280 nm is also split into two peaks at ∼262 and ∼274 nm. Similar changes were observed previously in the CD spectrum of Calf-Thymus DNA upon Dox binding by Agudelo et al. (DNA:Dox; 5:1).^25^ It was further commented that, while the change in intensity is associated with base destacking, the shift in the peaks is due to B-A transition^.25^. The low-intensity, negative peak at around 295 nm in the spectrum of DDD-Dox indicates unwinding of the dsDNA.^26^ Surprisingly, binding of Dox to DDD8 did not lead to any such change in the CD spectrum, which remains fairly similar to that of DDD8 in the absence of a small molecule. Overall, Dox binding appears to leave the structure of DDD8 nearly unchanged while it significantly disrupts that of DDD.

**Figure 3D** displays the changes in the CD spectra of DDD and DDD8 upon Hoe binding. No significant difference was observed upon binding of Hoe to DDD or to DDD8. Only a very slight intensity change at ∼250 nm with a slight shift to a higher wavelength (i.e., about 3 nm). An induced band formation was also observed between ∼300 and 400 nm in both DDD-Hoe and DDD8-Hoe complexes.

Thermal denaturation experiments of DDD and DDD8 were also performed in the presence of small molecules **(Figure 4 and 5, and Table 3)** using UV-Vis spectroscopy. Except for EtBr, all the small molecules were found to stabilize both DDD and DDD8. Under these conditions, EtBr was found to slightly destabilize both DDD and DDD8. The destabilizing effect of EtBr, based on the environment and the sequence, was discussed in previous studies.^27,28^ Dox, Net, and Hoe tend to stabilize DDD8 slightly more than DDD. And, the minor groove binders increase the stability of the dsDNAs more than Dox does. Finally, fluorescence titrations were carried out to determine the effect of DNA methylation on the binding affinity of the small molecules to DNA. Association constants (K_a_) obtained for EtBr, Dox and Hoe are given in **Table 4**, and the representative titration curves are given in **Supplementary Figure S3**. We could not determine the association constant for DDD-Net and DDD8-Net due to insignificant change in fluorescence upon DNA binding, as also discussed by Wartell et al.^29^ Overall, all the small molecules were found to be binding the methylated DDD8 **(Supplementary Figure 3 B, D, F)** slightly better than DDD **(Supplementary Figure 3 A, C, E)**. EtBr shows the weakest association with dsDNA among other small molecules considered in this work (4.2 x 10^4^ M^-1^ and 6.7 x 10^4^ M^-1^ with DDD **(Supplementary Figure 3A)** and DDD8 **(Supplementary Figure 3B)**, respectively). DDD-Dox **(Supplementary Figure 3C)**, DDD8- Dox **(Supplementary Figure 3D)**, DDD-Hoe **(Supplementary Figure 3E)** and DDD8-Hoe **(Supplementary Figure 3F)** complexes form with a comparable association constant in the range 2.38-5.80 10^5^ M^-1^. We note that the association constant of DDD8-Dox **(Supplementary Figure 3D)** is roughly twice as large as that of DDD-Dox **(Supplementary Figure 3C)**, which corroborates the observations by Evison et al. regarding the higher selectivity and enhanced toxicity of Dox toward methylated DNA.^30^

**Figure 4.**
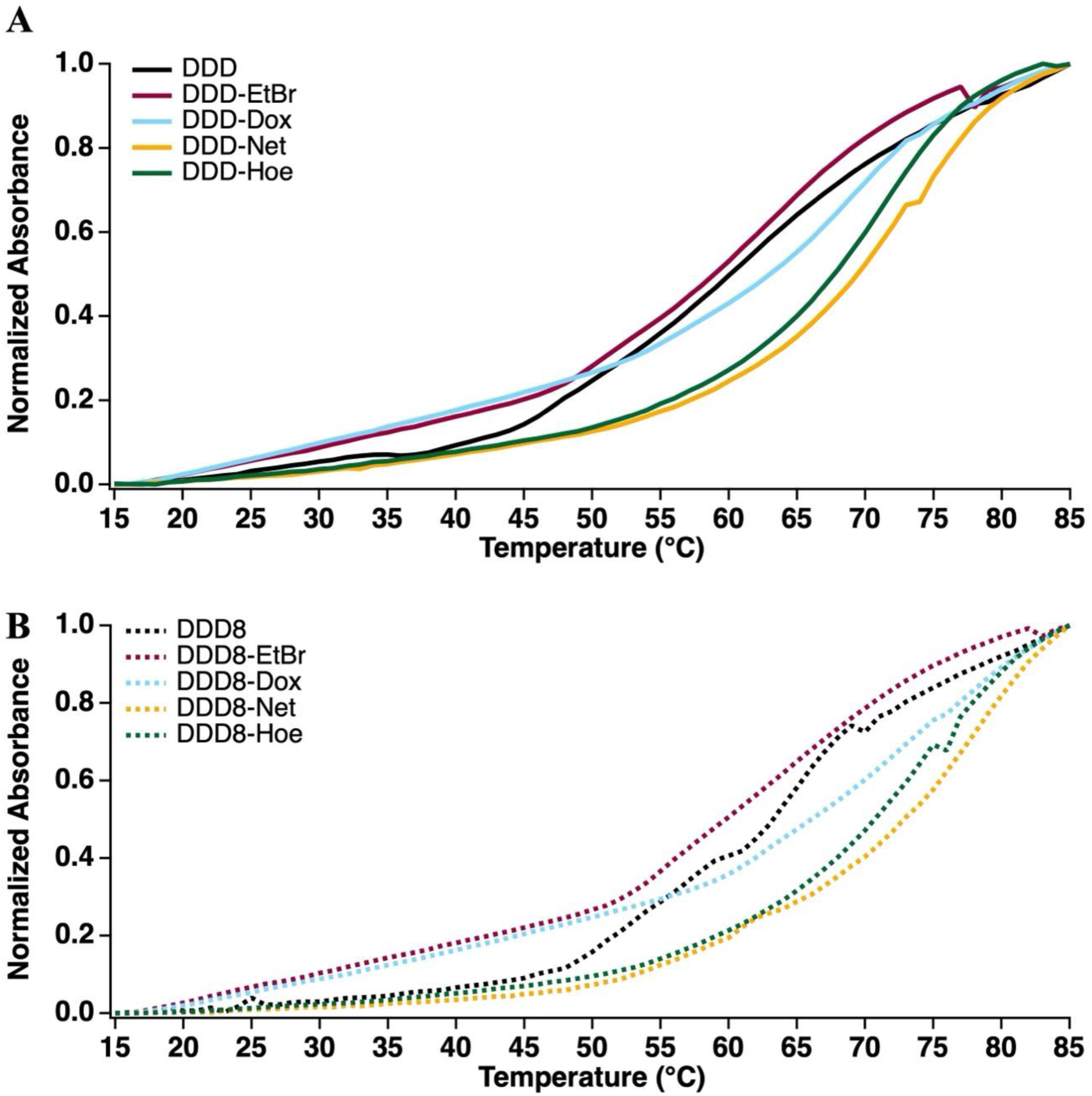
Average normalized thermal denaturation curves for DDD **(A)** and DDD8 **(B)** in the absence and presence of small molecules obtained by observing the UV-Vis absorbance changes at 260 nm with increasing temperature.

**Figure 5.**
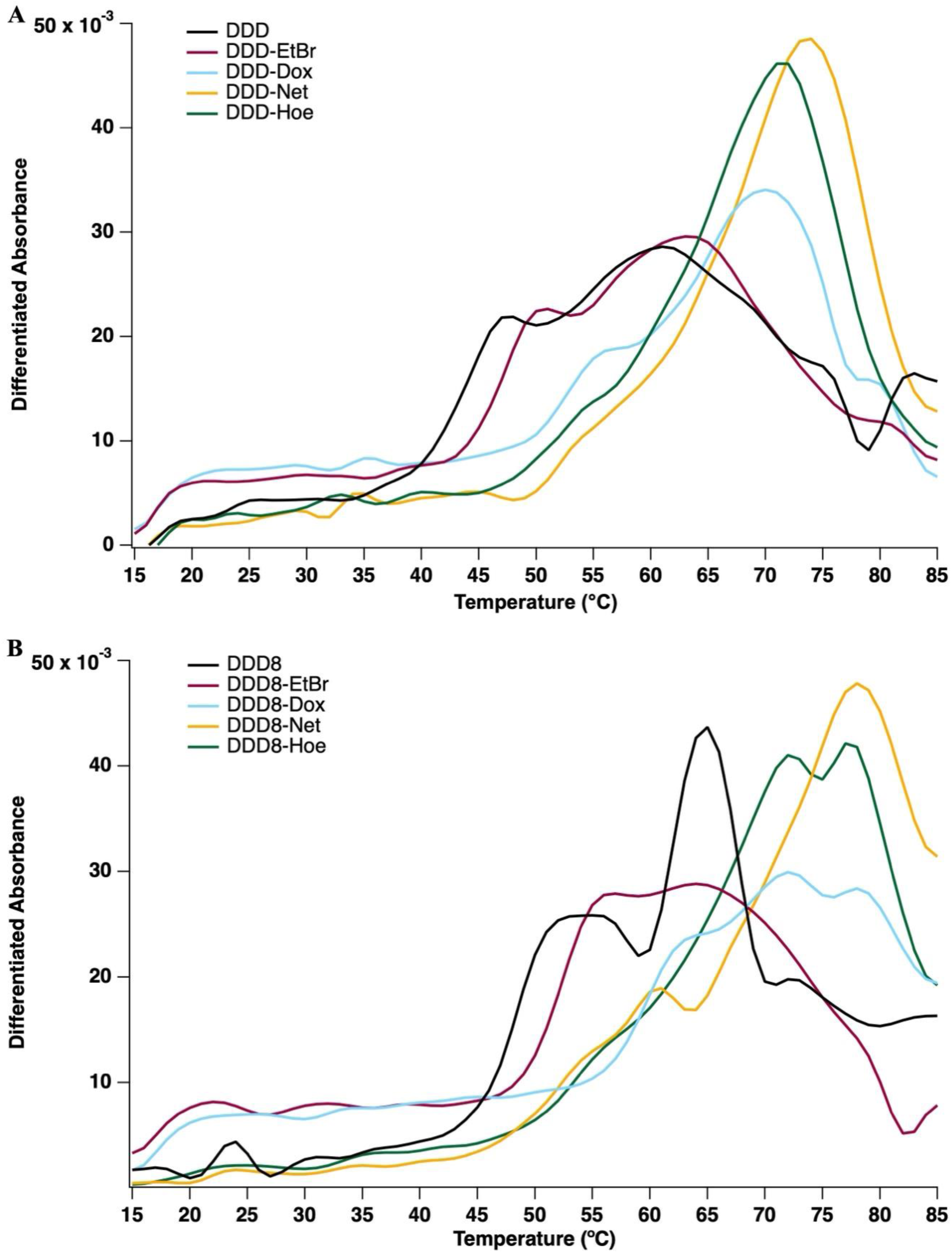
Differentiated thermal denaturation curves for DDD **_(A)_** and DDD8 **_(B)_** in the absence and presence of small molecules obtained by observing the UV-Vis absorbance changes at 260 nm with increasing temperature.

**Table 3.** Thermal denaturation temperatures (T_m_) of DDD, DDD8, and their complexes with small molecules obtained from the heating traces given in Figure 4.

| Sequence | $T_m$ (°C) | Sequence | $T_m$ (°C) |
| --- | --- | --- | --- |
| **DDD** | $60.0 \pm 0.5$ | **DDD8** | $63.5 \pm 0.1$ |
| **DDD-EtBr** | $59.0 \pm 0.1$ | **DDD8-EtBr** | $59.8 \pm 0.3$ |
| **DDD-Dox** | $63.3 \pm 0.3$ | **DDD8-Dox** | $66.3 \pm 0.3$ |
| **DDD-Net** | $69.5 \pm 0.5$ | **DDD8-Net** | $72.8 \pm 1.3$ |
| **DDD-Hoe** | $68.0 \pm 0.0$ | **DDD8-Hoe** | $70.8 \pm 0.3$ |

**Table 4.** Association constants (K_a_) obtained for small molecules with DDD and DDD8.

| Sequence | $K_a \cdot 10^5 \text{ (M}^{-1}\text{)}$ |
| --- | --- |
| DDD-EtBr | $0.42 \pm 0.02$ |
| DDD8-EtBr | $0.67 \pm 0.03$ |
| DDD-Dox | $2.38 \pm 0.12$ |
| DDD8-Dox | $5.80 \pm 0.59$ |
| DDD-Hoe | $3.60 \pm 0.36$ |
| DDD8-Hoe | $4.50 \pm 0.19$ |

The striking difference between DDD-Dox and DDD8-Dox CD spectra makes these two systems a valuable platform to further gain insights into the structural features of these complexes. In this direction, molecular dynamics (MD) simulations were performed for DDD and DDD8, in their apo form as well as complexed with Dox at different intercalation sites. A total of 26 systems were simulated in explicit water with two 750 ns-long MDs each to increase convergence of the results. Root mean square deviation time series for DNA and Dox are plotted in **Supplementary Figures S4** and **S5**, respectively. RMSD analysis indicates that the structures rapidly reach equilibrium, leading us to perform all further analysis on the last 500 ns of each simulation run. Figure 6 summarizes the findings obtained from the analysis of modelling data while additional data on structural descriptors of the dsDNA can be found in **Supplementary Figures S6-12.**

**Figure 6.**
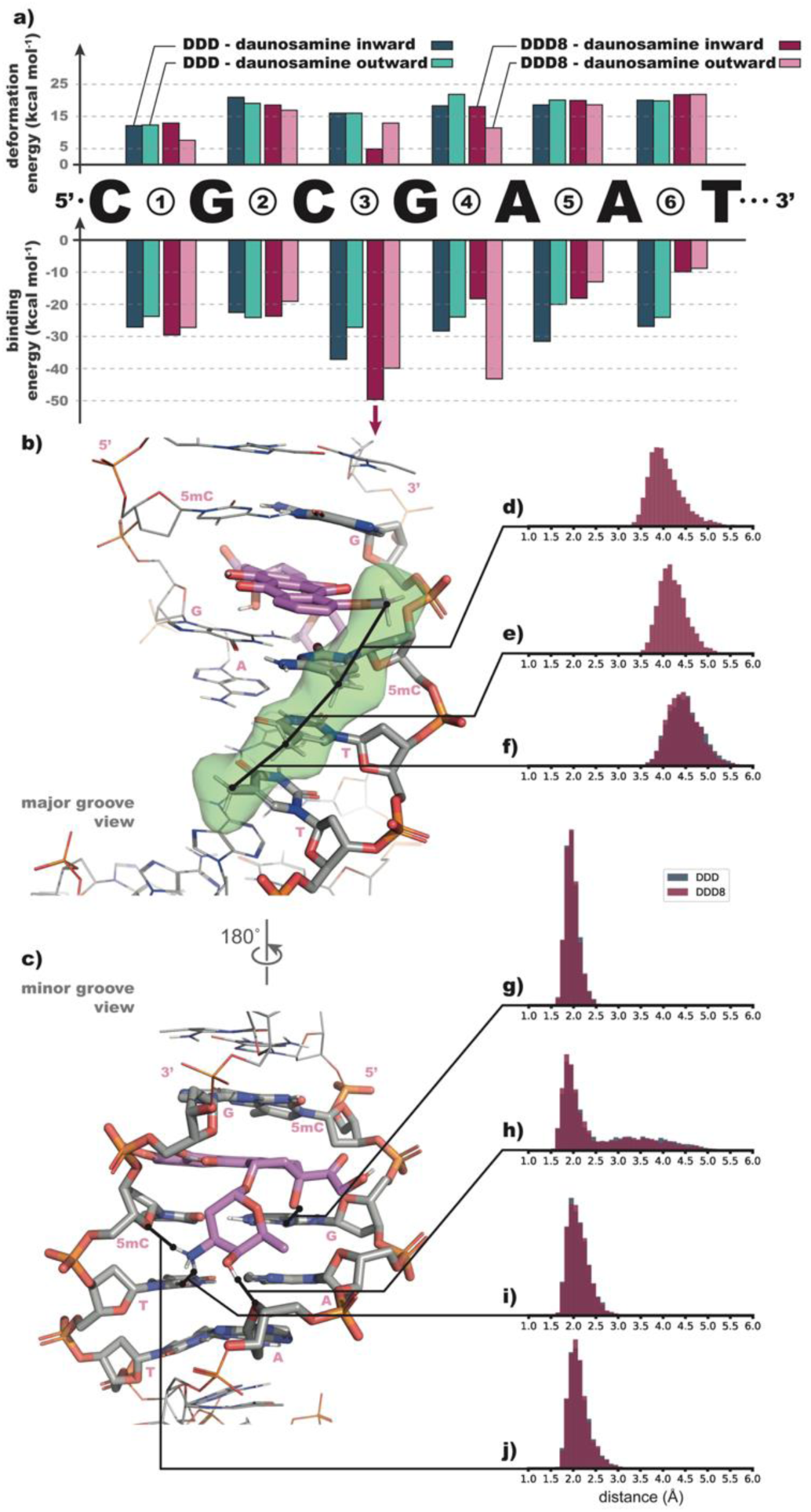
Comparative analysis of intercalation modes for DDD-Dox and DDD8-Dox. **(A)** Binding energy (including deformation and desolvation contributions) and deformation energy in kcal mol^-1^ as calculated for each of the 24 DNA-Dox complexes considered in this work. **(B-C)** Side view (major and minor groove, respectively) of a representative structure for the best binding mode of DDD8-Dox showing the main interactions between DNA and Dox. DNA and Dox carbon atoms are colored in grey and purple, respectively. The solvent accessible area around selected methyl groups in the major groove is represented in green color. **(D-J)** Distribution of selected inter atomic distances over the simulation time. **F and J** show data for DDD-Dox and DDD8-Dox (data significantly overlap), while **D** and **E** only show data for DDD8-Dox only since there is no methyl group on cytosines of DDD.

The data reported here relate to complexes in which the daunosamine group of Dox is placed in the minor groove of the dsDNA. Complexes with a daunosamine group in the major groove were also considered but led to highly unstable structures and were further disregarded (data not shown).^31^ Considering the symmetry of the DDD and DDD8 sequences, six intercalation sites with two orientation of the daunosamine group of Dox, either pointing towards the center of the sequence or towards its end, which are further referred to as “inward” and “outward” orientations, respectively. The binding energy at each intercalation site is reported in **Figure 6A** and includes desolvation energy within the MMGBSA model as well as deformation energy contributions. The deformation energy is also reported separately in the figure to emphasize the effect of Dox intercalation on the structure of the dsDNA. In both DDD and DDD8, Dox appears to favor intercalation site 3 (i.e., the second CpG step) with the daunosamine group pointing towards the center of the sequence. The binding energy of Dox with DDD8 at site 3 (about -50 kcal mol^-1^) is 13 kcal mol^-1^ more favourable than with DDD (about -37 kcal mol^-1^) and matches fairly well the higher affinity of Dox towards DDD8 observed experimentally **(Table 4)**. Besides the best intercalation mode, Dox shows a significant affinity towards sites 4, 5, and 6 of DDD with daunosamine inward and binding energies of -28, -30, and -27 kcal mol^-1^, respectively. The low selectivity of Dox with respect to AT and CG sites was also observed by Airoldi et al.^32^ In DDD8, two other intercalation modes have a strong affinity at sites 3 and 4 with the daunosamine pointing in the other direction (outward) and energies of -40 and -43 kcal mol^-1^, respectively. Analysis of RMSD time series of Dox **(Supplementary Figure S5)** shows that intercalation at sites 3, 4, 5, and 6 with daunosamine inward produce complexes that are significantly more stable than their counterpart with daunosamine outward. This observation is in line with the available experimental structures of dsDNA-Dox complexes showing only the inward orientation of the daunosamine group.^33^ Disregarding the complexes with daunosamine pointing outward due to their lower stability, the best binding mode in DDD8 (i.e., at site 3) appears to be more than 20 kcal mol^-1^ more favourable than any other site.

Analysis of structural parameters for DDD and DDD8 in their apo or complexed forms revealed some differences regarding the impact of the intercalator on key geometrical features. The major groove width **(Supplementary Figure S6)** is left nearly unaffected by the intercalation. Intercalation at site 4 with daunosamine inward and at sites 2, 3, and 4 with daunosamine outward tend to increase the minor groove’s width, producing an additional peak in the distribution plotted in **Supplementary Figure S7**. Analysis of other geometrical parameters **(Supplementary Figures S8-12)** show that all intercalation sites have an impact on surrounding DNA bases, either locally or up to bases located one step away from the intercalator. Although our analysis of the present simulations only shows deviations from B-form, it is in line with the partial B to A transition observed by Agudelo et al. upon Dox intercalation.^34^ Site 3 with the daunosamine group of Dox pointing inward is a notable exception and shows little to no disrupting effect on the dsDNA structure. Both DDD and DDD8 behave similarly upon intercalation at this site and experience stiffening of their structure with a significant decrease in the width of the corresponding distribution of χ angle, x-displacement, phase angle, and slide parameters.

Carvalho et al. studied the effect of methylation on DDD by considering 5mC at positions 3, 9, and both 3 and 9.^35^ The authors concluded that methylation at position 3 changes the hydrophobicity of sequence and alters its hydration pattern. To accommodate this change, the dsDNA unwinds, which causes an increase in twist. However, methylation at position 9 does not affect the structure of DNA because the methyl group of 5mC and that of the two thymidines (at position 7 and 8) are on the same strand and form a stable chain of three hydrophobic groups, which eventually contributes to the stability of the dsDNA. A closer look at the structure of the complex formed by the intercalation of Dox in DDD8 at site 3 with daunosamine pointing inward is depicted in **Figures 6B** and **6C**. Key interatomic distances characterizing relevant intermolecular interactions were monitored along the dynamics of the complex and their distribution over the last 500 ns of each simulation run is given in **Figures 6D-J** for DDD and DDD8. In DDD8, the methoxy groups of Dox interacts with the methyl group of 5mC (i.e. at position 9) and thereby contributes to a chain of four methyl groups forming a stable hydrophobic cluster with the two thymidines (i.e., at positions 7 and 8) at the centre of the sequence **(Figures 6B** and **6D-H)**. Such interaction is not possible in DDD since the cytosines lack the methyl group and correspondingly the data in **Figures 6D** and **6E** only report the distribution for DDD8. The interactions between Dox and dsDNA in the minor groove are dominated by polar interactions. Dox interacts with the guanosine at the intercalation site **(Figure 6G)** with a highly stable hydrogen bond, and with the sugar of the second adenosine **(Figure 6H)** with a slightly looser hydrogen bond. Dox also interacts with the other strand via its amine group by forming strong hydrogen bonds with the sugar of the cytosine at the intercalation site **(Figure 6I)** and with the nitrogen of the previous thymine base **(Figure 6J)**. The comparison of the distribution of these critical distances between DDD and DDD8 **(Figures 6F-J)** shows a nearly identical interaction pattern and are in line with previous observations.^36^ The only difference between DDD-Dox and DDD8-Dox complexes is thus the chain of hydrophobic interactions that is only present in the major groove of DDD8.

Considering the results obtained from the experimental and modelling parts of the present study, we postulate that Dox is highly selective towards a single intercalation site in DDD8 (site 3 with daunosamine inward) while, as shown by Airoldi et al.^32^, it is significantly less selective in DDD. The modelling data indicate that intercalation at site 3 does not affect the structure of DNA, which matches well with the experimental observation showing that the CD spectrum of DDD8 and DDD8**-**Dox are very similar. This further suggests that, in the case of DDD8, the intercalation occurs solely at site 3 with daunosamine pointing inward. On the other hand, the lower selectivity of Dox for a specific site in DDD lets us speculate that the intercalation can occur at other sites than site 3, which would affect the structure of the dsDNA. The CD spectrum obtained for DDD-Dox is therefore likely to result from a mixture of complexes with Dox intercalated at different sites.

## CONCLUSIONS

The role of DNA methylation on DNA structure and stability was investigated by systematically changing the number and the placement of 5-methylcytidines in a model Dickerson dodecamer (DDD). Our results clearly demonstrate that DNA methylation increases the stability and changes the conformation of DNA only with slight deviations from the B-form conformation. Yet even slight alterations have a significant effect on DNA-small molecules interactions. All the small molecules investigated in the present work (i.e., EtBr, Dox, Net, and Hoe) were found to bind to DDD8, (i.e., DDD with eight d5mCs) with a slightly higher affinity compared to DDD. On the other hand, based on the CD spectral changes, they appear to alter the structure of DDD more than that of DDD8. Especially, Dox was found to be altering DDD structure significantly in a similar way to its alteration of Calf-Thymus DNA,^25^ while no such dramatic conformational change was observed upon binding to DDD8. Further investigations of DDD-Dox and DDD8-Dox complexes via molecular dynamics simulations shed light onto the rational of the difference in these structural changes. Dox can form a hydrophobic cluster involving a chain of four methyl groups upon binding to DDD8, while such interaction is not possible upon binding to non-methylated DDD. The formation of the hydrophobic cluster significantly increases the binding selectivity of Dox to a specific intercalation site in DDD8 (site 3; 5’- dTdTd5mCdG-3’). We believe that creating such hydrophobic clusters using small molecules might play a role in regulating gene expression selectively in specific hypermethylated sites. In brief, the results rationalize the effect of methylation on DNA-small molecule interactions and pave the way to new avenues in the design of novel compounds targeting specifically hypermethylated or hypomethylated sites in chemotherapy.

## LIST OF ABBREVIATIONS

d5mC: the methylation of cytosines at the 5^th^ position
DNA: deoxyribonucleic acid
DDD: Dickerson dodecamer
EtBr: ethidium bromide
Dox: doxorubicin
Net: netropsin
Hoe: hoechst 33258
dC: cytidine
dsDNA: double stranded DNA
HMT: histone methyltransferase
DNMT: DNA methyltransferase
CD: circular dichroism
UV-Vis: UV-visible
nab: nucleic acid builder
GBSA: generalized Born/surface area
T_m_: thermal denaturation temperature
K_a_: association constant
MD: molecular dynamics

## ACKNOWLEDGEMENTS

We would like to thank The Scientific and Technological Research Council (TÜBİTAK-118Z339) of TÜRKİYE and Middle East Technical University Scientific Research Projects Coordination Unit (METU BAP) (AGEP-103-2019-10327) for financial support. We also would like to thank Prof. Dr. Ahmet Önal’s group (Middle East Technical University, Turkey) for providing the opportunity for fluorescence spectroscopy measurements. The numerical calculations reported in this paper were partially performed at TÜBİTAK ULAKBİM, High Performance and Grid Computing Center (TRUBA resources). We are also grateful to Dr. Antonio Monari (Université de Lorraine, France) for valuable discussions.

