## Supplementary Information for "The effect of DNA methylation on DNA structure and DNA-small molecule interactions"

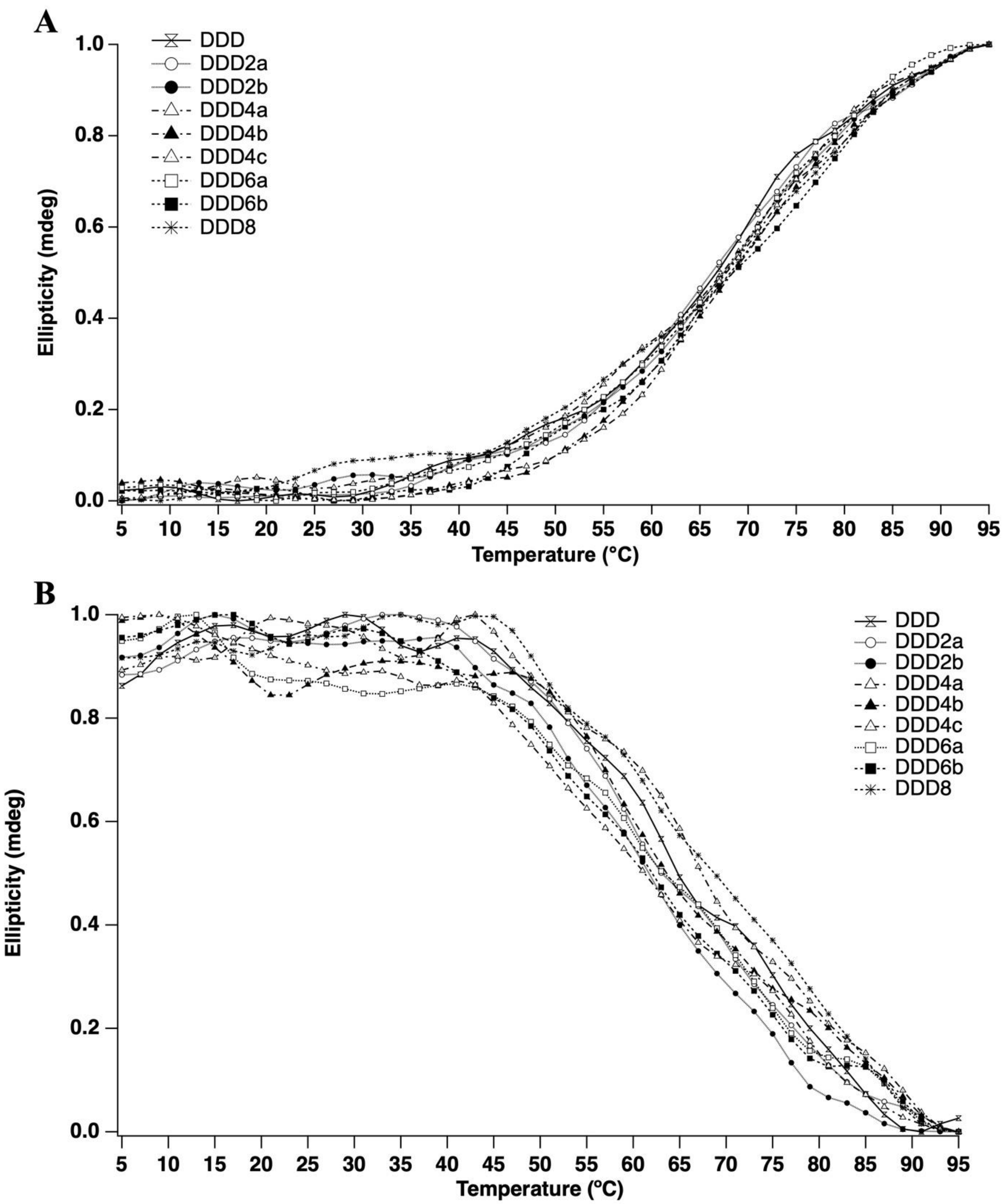


**Supplementary Figure S1.** Normalized thermal denaturation curves for DDD and d5mC substituted DDDs obtained by observing the CD ellipticity changes at 250 nm **(A)** and at 279 nm **(B)** with increasing temperature.

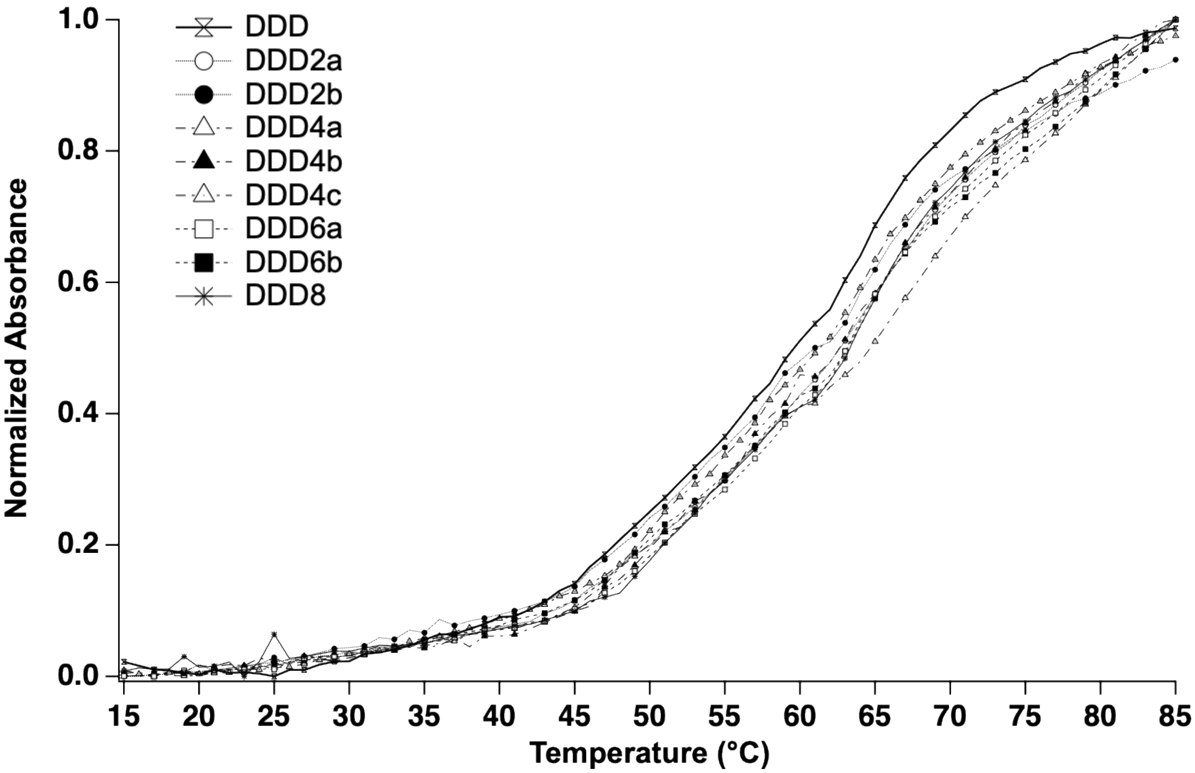


**Supplementary Figure S2.** Normalized thermal denaturation curves for DDD and d5mC substituted DDDs obtained by observing the UV-Vis absorbance changes at 260 nm with increasing temperature.

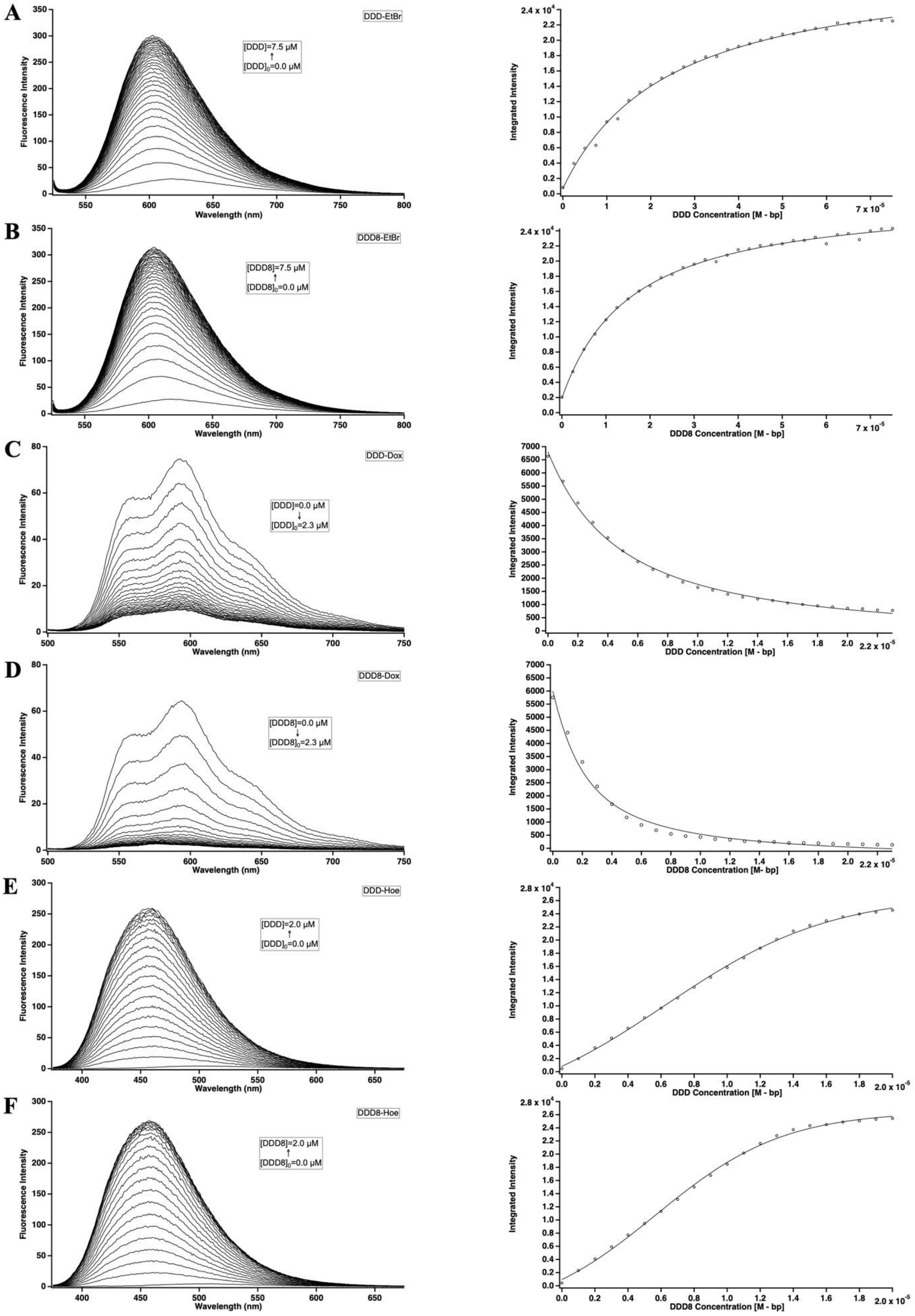


**Supplementary Figure S3.** Fluorescence intensity vs wavelength spectra of DNA solutions which were titrated with the solutions including 1 μM of small molecule and the plots of integrated fluorescence intensity vs DNA concentration.

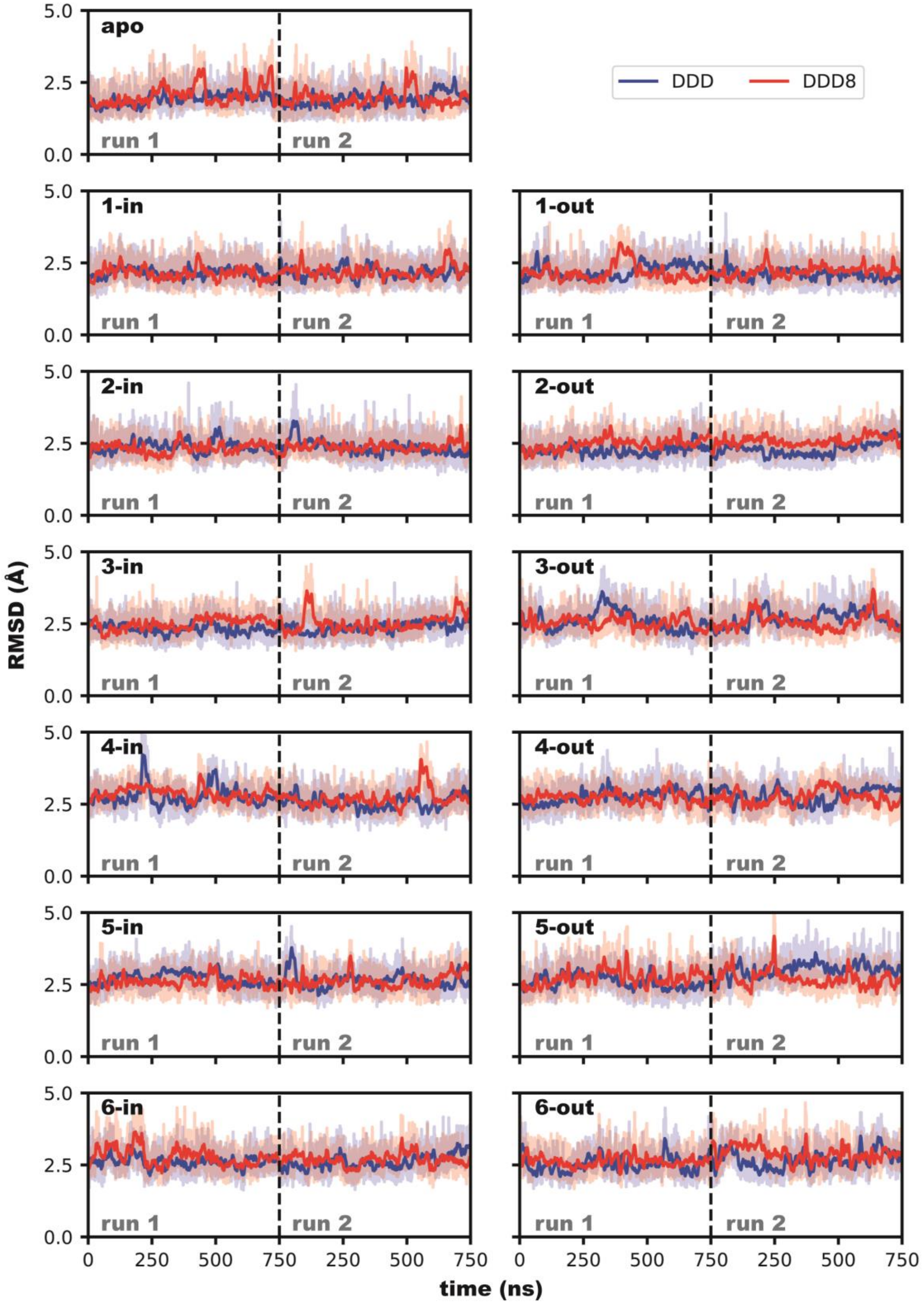


**Suplementary Figure S4.** RMSD of DNA vs time for runs of each MD system.

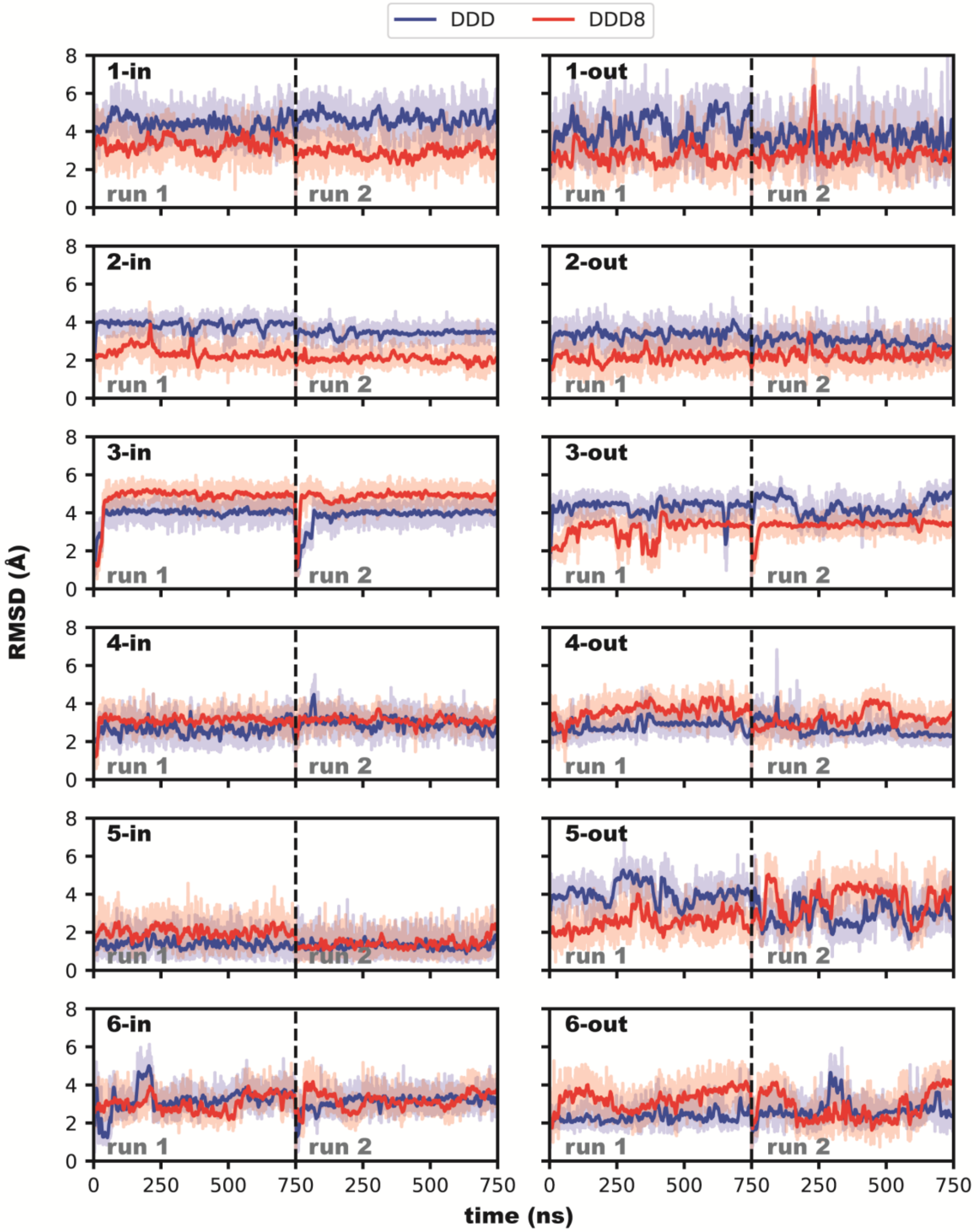


**Supplementary Figure S5.** RMSD of Dox vs time for runs of each MD system.

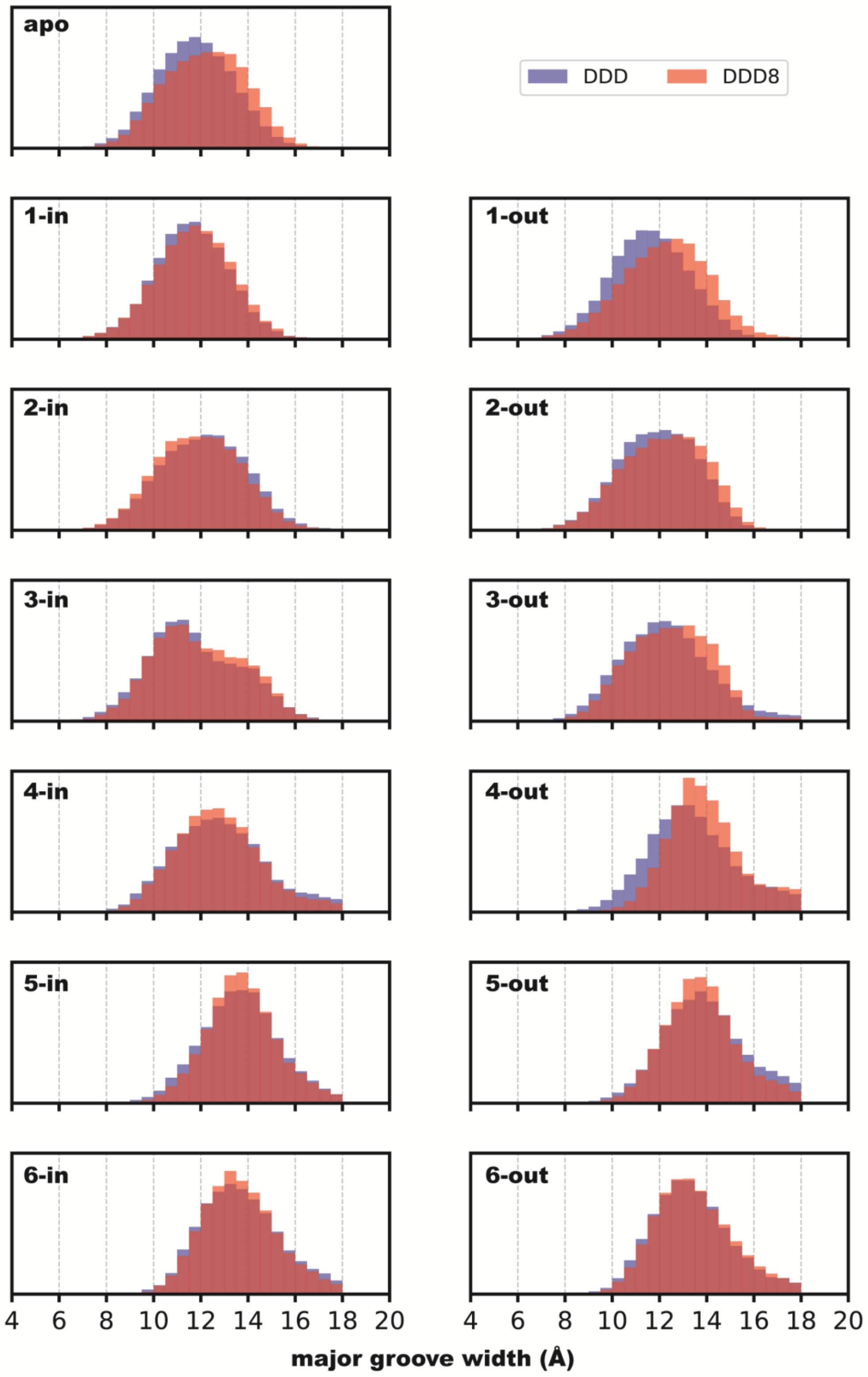


**Supplementary Figure S6.** Major groove width distributions for each MD system of DDD8 and DDD.

DDD.

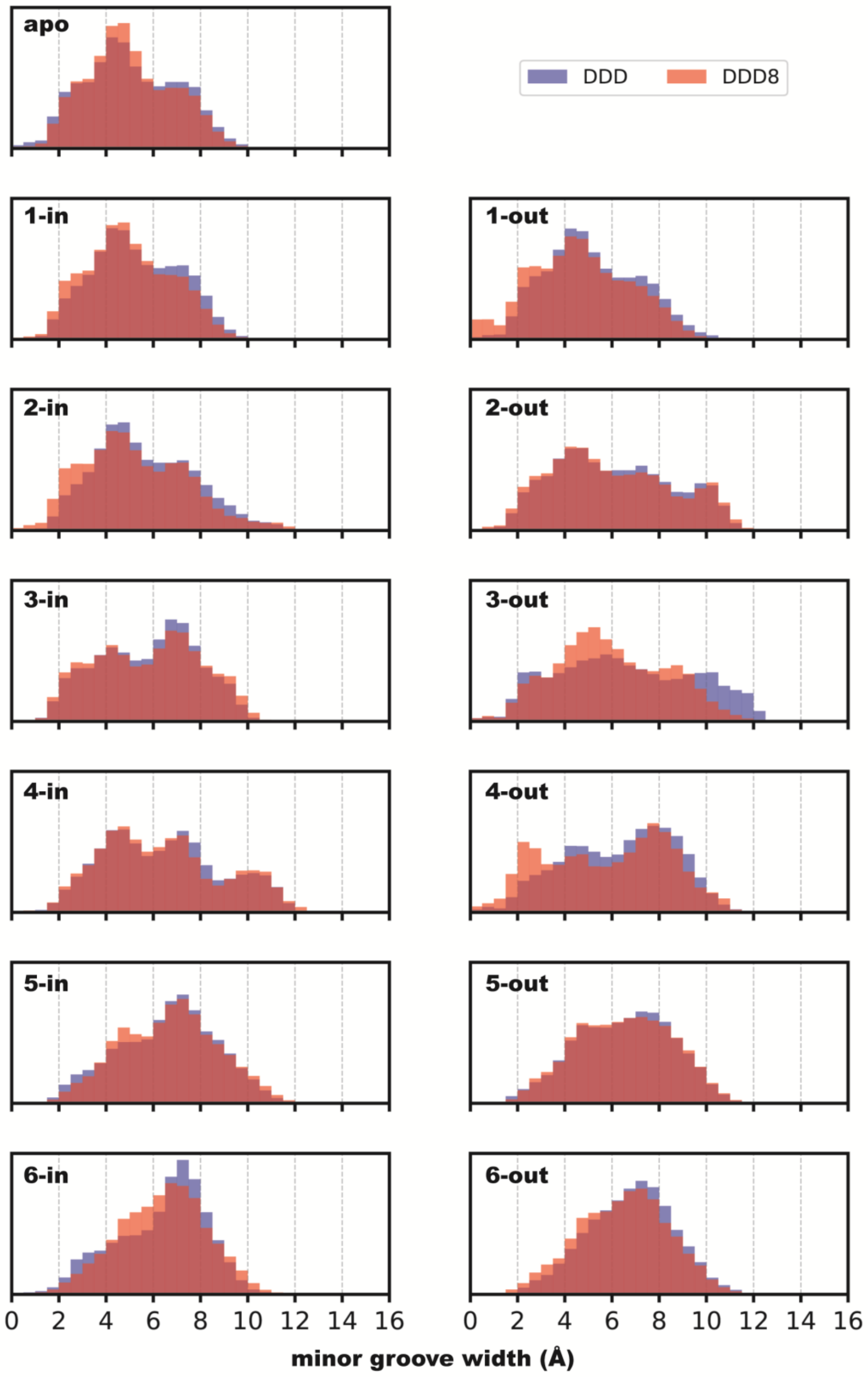


**Supplementary Figure S7**. Minor groove width distributions for each MD system of DDD8 and DDD.

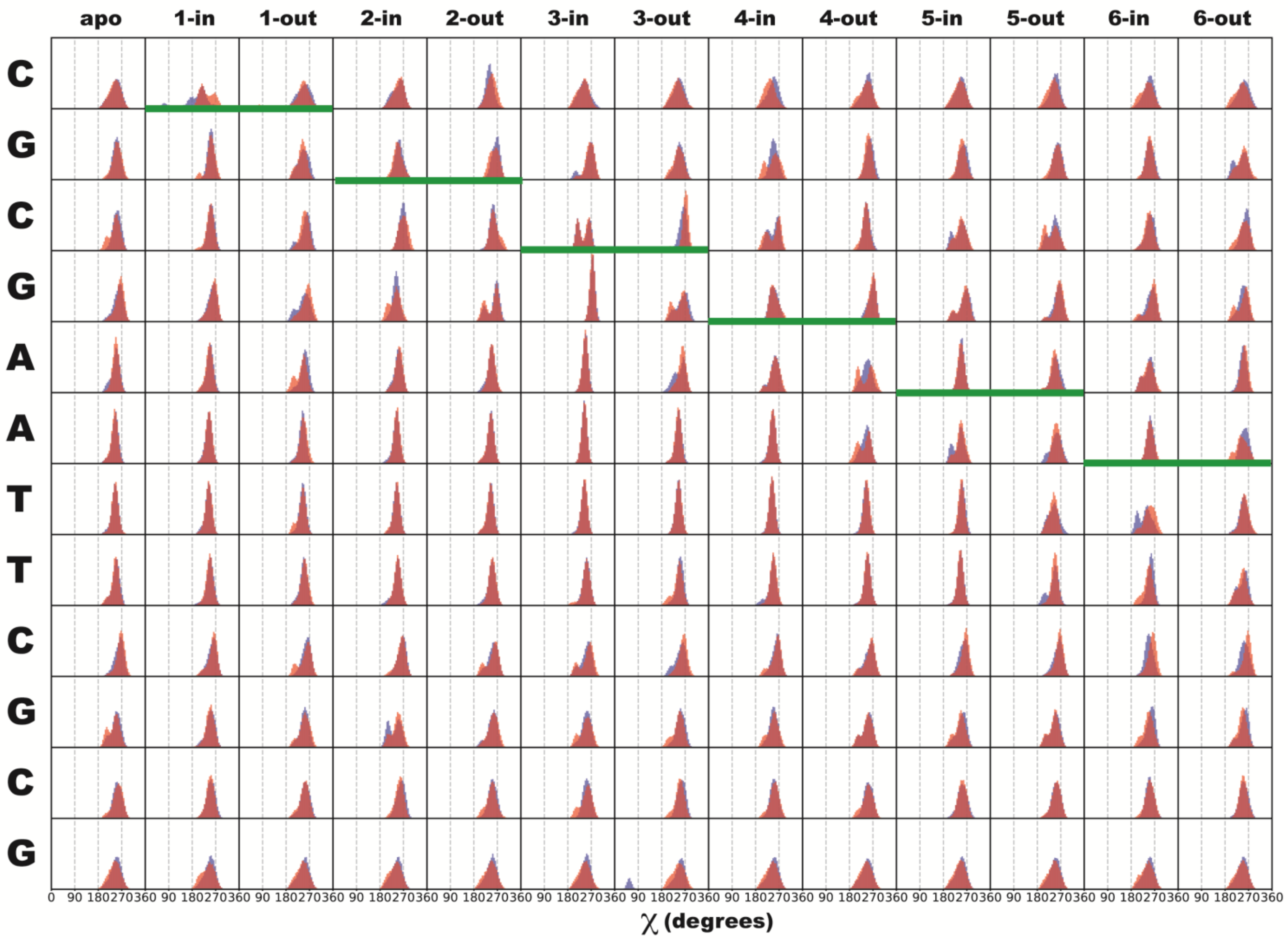


**Supplementary Figure S8.** Chi parameter distributions for base pairs of each MD system of DDD and DDD8. Each column refers to different systems and each row refers different base pair steps. Green thick lines indicate the position of intercalation in each system.

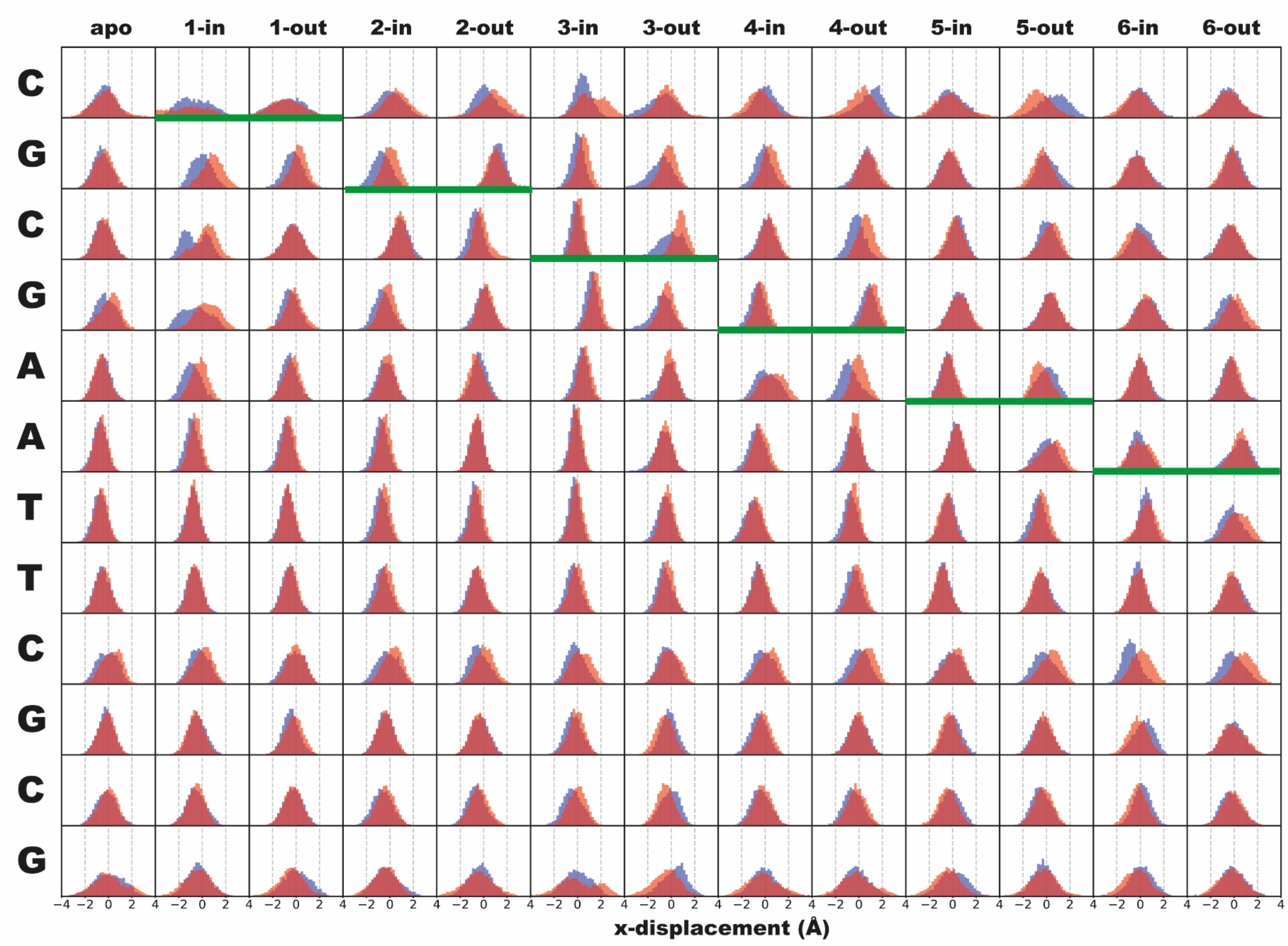


**Supplementary Figure S9.** X displacement parameter distributions for base pairs of each MD system of DDD and DDD8. Each column refers to different systems and each row refers different base pair steps. Green thick lines indicate the position of intercalation in each system.

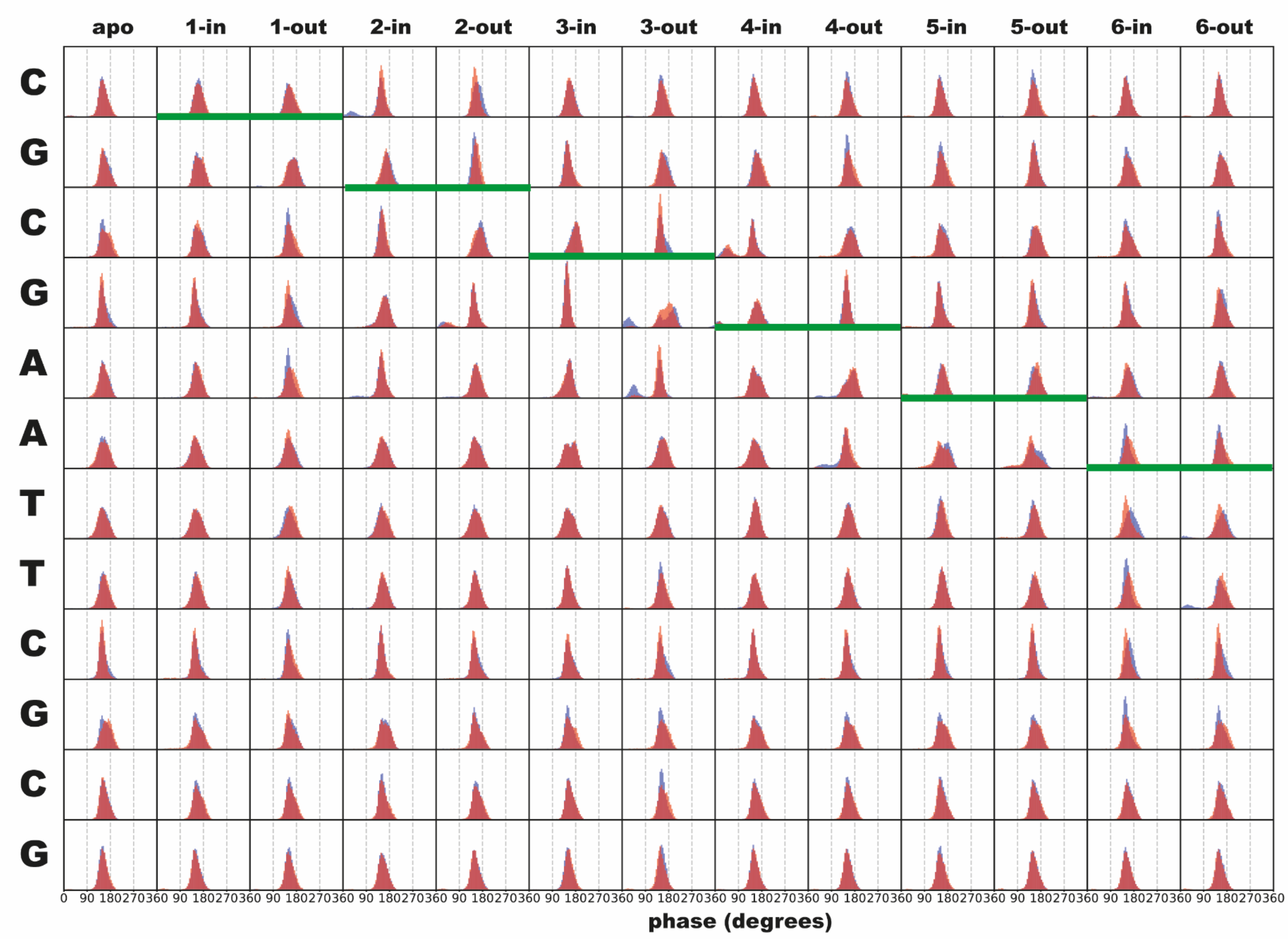


**Supplementary Figure S10**. Phase parameter distributions for base pairs of each MD system of DDD and DDD8. Each column refers to different systems and each row refers different base pair steps. Green thick lines indicate the position of intercalation in each system.

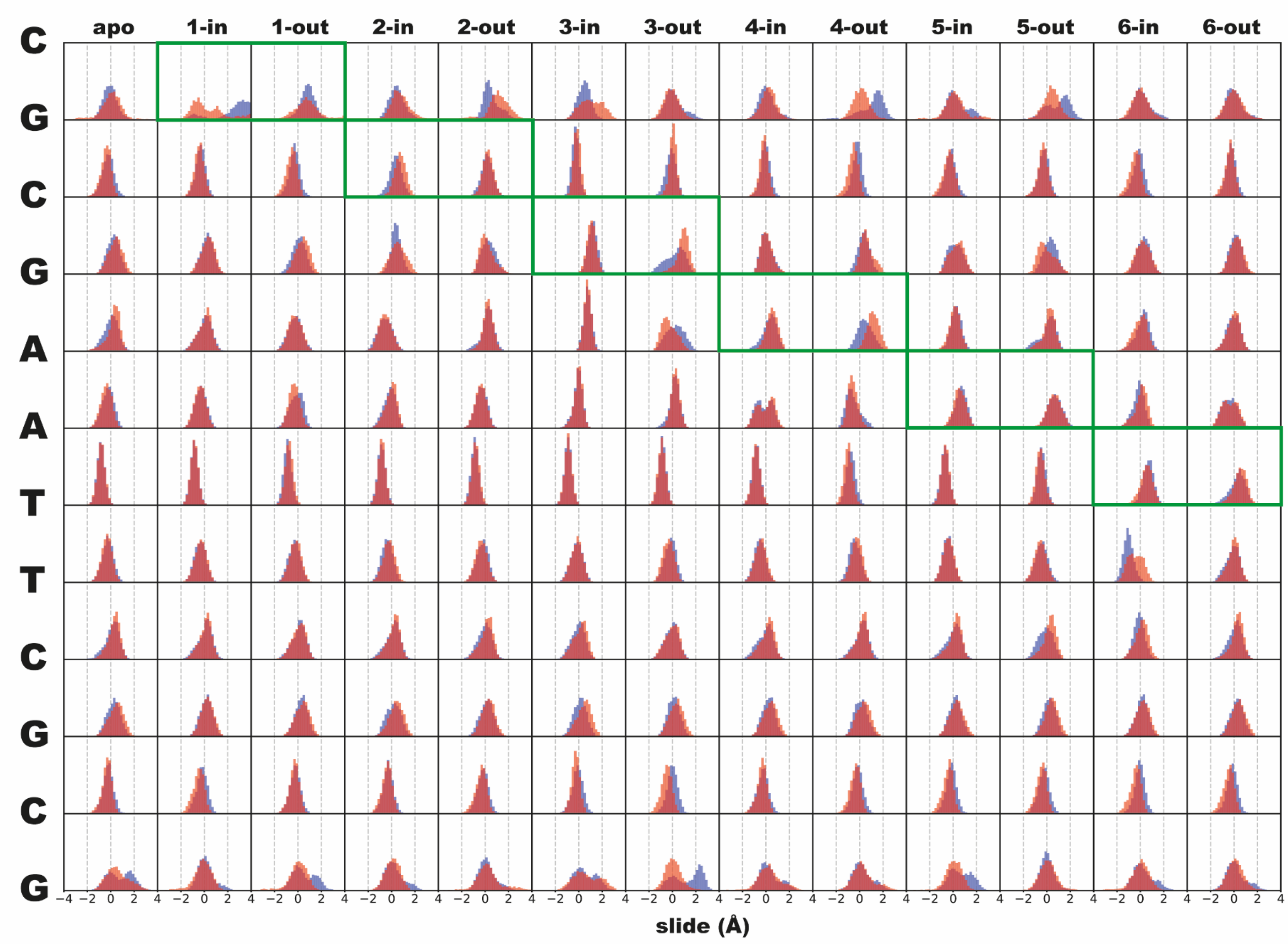


**Supplementary Figure S11**. Slide parameter distributions for successive base pairs of each MD system of DDD and DDD8. Each column refers to different systems and each line refers different base pair steps. Green rectangles indicate the position of intercalation in each system.

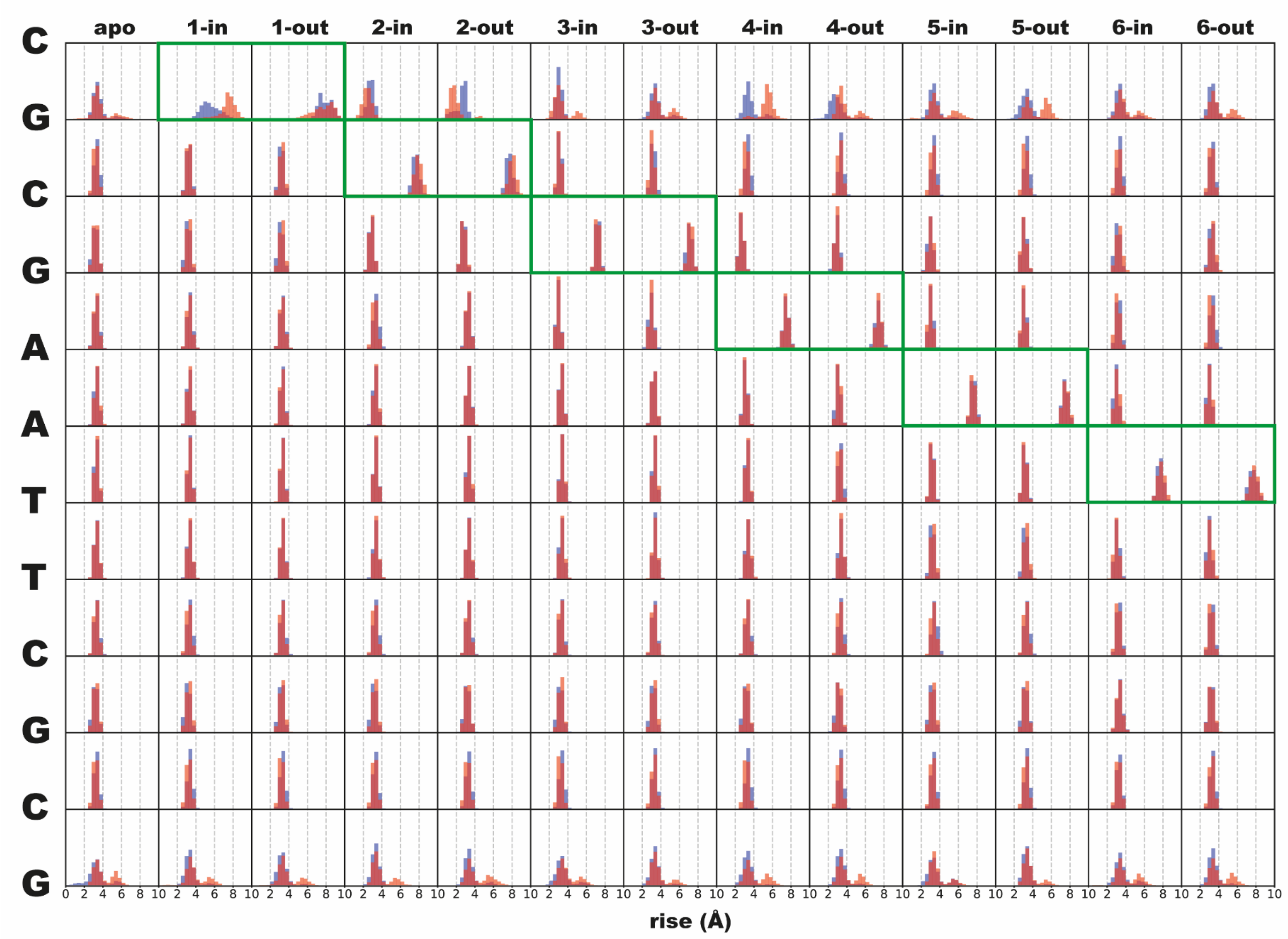


**Supplementary Figure S12**. Rise parameter distributions for successive base pairs of each MD system of DDD and DDD8.Each column refers to different systems and each line refer different base pair step. Green rectangles indicate the position of intercalation in each system.
